# A chemoproteomic portrait of the oncometabolite fumarate

**DOI:** 10.1101/285759

**Authors:** Rhushikesh A. Kulkarni, Daniel W. Bak, Darmood Wei, Sarah E. Bergholtz, Chloe A. Briney, Jonathan H. Shrimp, Abigail L. Thorpe, Arissa Bavari, Aktan Alpsoy, Michaella Levy, Laurence Florens, Michael P. Washburn, Emily C. Dykhuizen, Norma Frizzell, Eranthie Weerapana, W. Marston Linehan, Jordan L. Meier

**Affiliations:** Chemical Biology Laboratory, Center for Cancer Research, National Cancer Institute, National Institutes of Health, Frederick, MD, 21702, USA.; Department of Chemistry, Boston College, Chestnut Hill MA, 02467, USA.; Urologic Oncology Branch, Center for Cancer Research, National Cancer Institute, National Institutes of Health, Bethesda, MD, 20817, USA.; Department of Medicinal Chemistry and Molecular Pharmacology, College of Pharmacy, Purdue University, West Lafayette IN, 47906, USA.; Stowers Institute for Medical Research, Kansas City, MO, 64110, USA; Department of Pathology and Laboratory Medicine, University of Kansas Medical Center, Kansas City, KS, 66160, USA; Department of Pharmacology, Physiology and Neuroscience, School of Medicine, University of South Carolina, Columbia, SC, 29209, USA.

## Abstract

Hereditary cancer disorders often provide an important window into novel mechanisms supporting tumor growth and survival. Understanding these mechanisms and developing biomarkers to identify their presence thus represents a vital goal. Towards this goal, here we report a chemoproteomic map of the covalent targets of fumarate, an oncometabolite whose accumulation marks the genetic cancer predisposition syndrome hereditary leiomyomatosis and renal cell carcinoma (HLRCC). First, we validate the ability of known and novel chemoproteomic probes to report on fumarate reactivity in vitro. Next, we apply these probes in concert with LC-MS/MS to identify cysteine residues sensitive to either fumarate treatment or fumarate hydratase (*FH*) mutation in untransformed and human HLRCC cell models, respectively. Mining this data to understand the structural determinants of fumarate reactivity reveals an unexpected anti-correlation with nucleophilicity, and the discovery of a novel influence of pH on fumarate-cysteine interactions. Finally, we show that many fumarate-sensitive and *FH*-regulated cysteines are found in functional protein domains, and perform mechanistic studies of a fumarate-sensitive cysteine in SMARCC1 that lies at a key protein-protein interface in the SWI-SNF tumor suppressor complex. Our studies provide a powerful resource for understanding the influence of fumarate on reactive cysteine residues, and lay the foundation for future efforts to exploit this distinct aspect of oncometabolism for cancer diagnosis and therapy.

## Introduction

A major finding of modern cancer genomics has been the unexpected discovery of driver mutations in primary metabolic enzymes.^1-5^ Many of these lesions cause the characteristic accumulation of “oncometabolites,” endogenous metabolites whose accretion can directly drive malignant transformation.^6^ For example, mutation of fumarate hydratase (*FH*) in the familial cancer susceptibility syndrome hereditary leiomyomatosis and renal cell carcinoma (HLRCC) leads to high levels of intracellular fumarate.^7-8^ Fumarate has been hypothesized to promote tumorigenesis both by reversibly inhibiting dioxygenases involved in epigenetic signaling,^9-14^ as well as by interacting with proteins covalently as an electrophile, forming the non-enzymatic posttranslational modification cysteine S-succination (Fig. 1).^15-16^ This latter mechanism is unique to fumarate, and has been proposed to contribute to the distinct tissue selectivity, gene expression profiles, and clinical outcomes observed in HLRCC relative to other oncometabolite-driven cancers.^17-18^ Consistent with a functional role, recent studies have found that S-succination of Keap1 can activate NRF2-mediated transcription in HLRCC.^19^ Furthermore, global immunohistochemical staining of S-succination has been applied to assess stage and progression of FH-deficient tumors,^20^ suggesting the utility of this modification as a biomarker.

**Figure 1.**
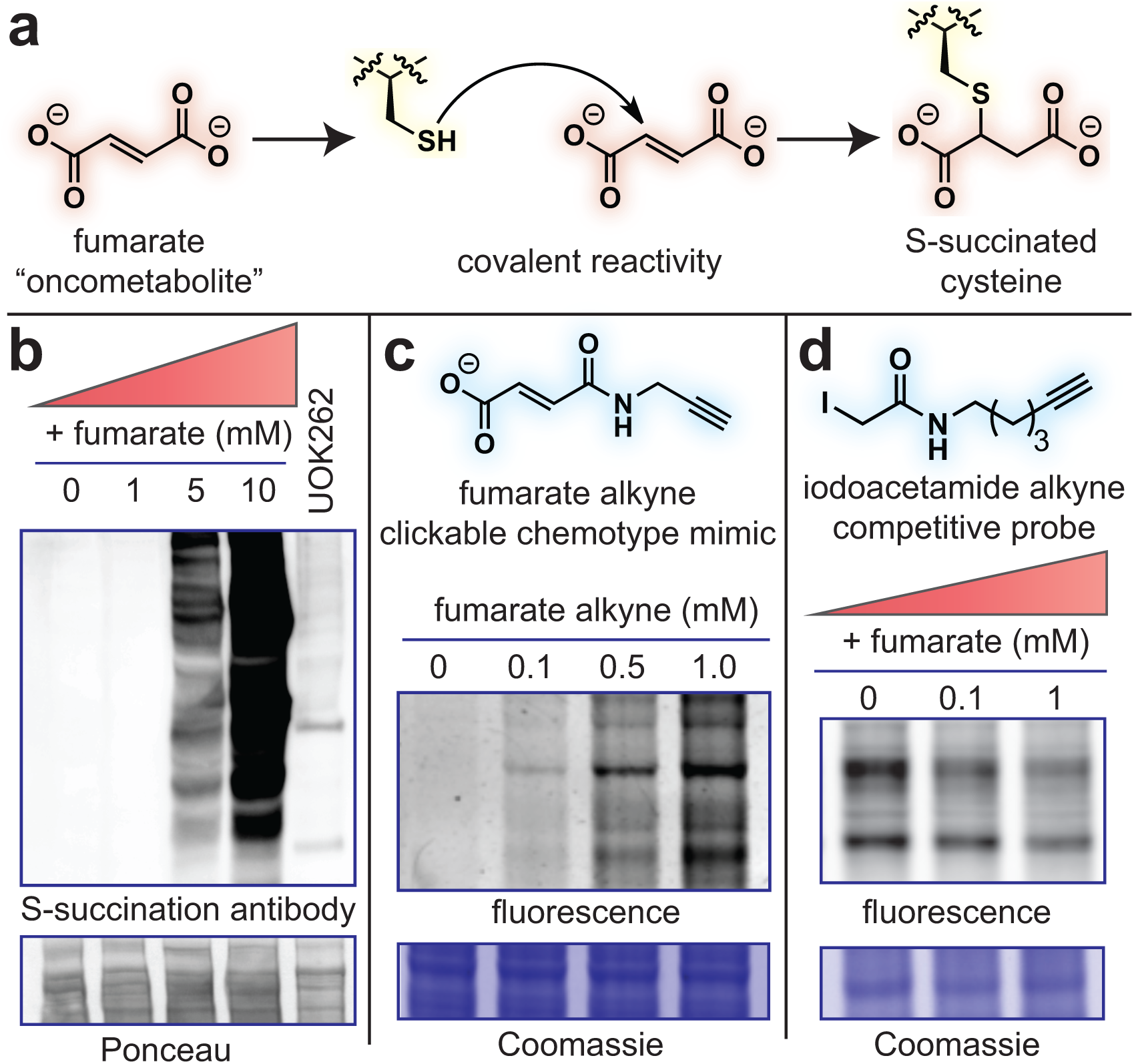
Fumarate is a covalent oncometabolite. (a) Covalent labeling of cysteine residues by fumarate yields the PTM S-succination. (b) S-succinated Cys immunoblotting establishes the physiological range of fumarate required for covalent protein labeling lies in the millimolar range. HEK-293 proteomes were treated with fumarate (0, 1, 5, 10 mM) for 15 h prior to western blotting. (c) Fumarate alkyne (FA-alkyne) can be used to visualize reactivity of the fumarate chemotype. HEK-293 proteomes were treated with FA-alkyne (0, 0.1, 0.5, 1 mM) for 15 h prior to click chemistry. (d) Iodoacetamide alkyne (IA-alkyne) can be used as a competitive probe of covalent fumarate labeling (HEK-293; 15 h pre-incubation with fumarate, then 100 μM IA-alkyne for 1 h followed by desalting and click chemistry).

Despite its potential relevance to HLRCC pathology, our overall understanding of fumarate’s covalent reactivity remains incomplete. Cysteine S-succination antibodies are not commercially available, and those developed have not yet proven useful for immunoprecipitation.^21^ This has limited our current knowledge of S-succination to proteins identified by candidate methods, such as Keap1,^19^ or whole proteome mass spectrometry,^22-25^ which is biased towards the identification of abundant proteins. In addition, neither of these approaches report on the extent of S-succination, impeding our ability to understand what structural interactions drive fumarate’s covalent reactivity, as well as whether these modifications significantly alter protein function. Thus, a better understanding of the global scope and stoichiometry of fumarate reactivity has the potential to provide new insights into HLRCC biology, as well as site-specific biomarkers for assessing tumor development and therapeutic response.

Towards this goal, here we report a chemoproteomic map of the covalent targets of the HLRCC oncometabolite fumarate. First, we establish the utility of chemoproteomic probes to report on concentration-dependent fumarate reactivity in vitro. Next, we apply these probes in combination with quantitative mass spectrometry to define the proteome-wide sensitivity of cysteine residues to either fumarate treatment or fumarate hydratase (*FH*) mutation. Analyzing this data to understand the molecular determinants of fumarate-sensitivity leads to the discovery of an unanticipated anti-correlation with cysteine nucleophilicity, and highlights a distinct impact of pH on the reactivity of this oncometabolite. Finally, we demonstrate that fumarate-sensitive and *FH*-regulated cysteines lie in many functional protein domains, and perform functional analyses of a fumarate-sensitive cysteine in SMARCC1, a member of the SWI-SNF tumor suppressor complex. Our studies provide a novel resource for understanding how fumarate reactivity impacts HLRCC biology, as well as an essential underpinning for diagnostic and therapeutic applications seeking to exploit this unique aspect of oncometabolism for clinical benefit.

## Results

### Comparative affinity-based profiling of physiological fumarate reactivity

Several recent studies have demonstrated the feasibility and power of applying reactivity-based chemoproteomics to characterize electrophilic drug targets.^26-29^ In order to extend these methods to the endogenous oncometabolite fumarate, we first sought to establish a physiological range for fumarate reactivity in complex proteomes. Several pieces of evidence imply that fumarate may exhibit relatively modest reactivity compared to other chemical electrophiles. Most relevantly, two recent analyses of the multiple sclerosis drug dimethyl fumarate (Tecfidera; DMF) found its metabolized product monomethyl fumarate (MMF) possesses limited thiol reactivity at micromolar concentrations.^30-31^ Theoretical calculations indicate this stems from MMF’s higher-lying LUMO, which increases the energetic barrier to covalent bond formation with nucleophilic cysteines.^31^ Of note, fumarate’s LUMO is even higher in energy than MMF (Fig. S1a), suggesting it may possess a distinct reactivity profile relative to Tecfidera. Further, it raises the question as to whether fumarate normally functions as a covalent metabolite, or does so only upon hyperaccumulation caused by pathophysiological stimuli such as *FH* mutation.

To explore this phenomenon, we treated cell lysates from human embryonic kidney (HEK-293) cells with increasing amounts of fumarate and analyzed proteins for covalent labeling using an S-succination antibody (Fig. 1b). An equal amount of proteome from the UOK262 HLRCC cell line (*FH*−/−) was used to compare levels of S-succination caused by *FH* mutation. Control studies verified the S-succination signal in these cells was dependent on *FH* mutational status (Fig. S1b). In contrast to its esterified analogues fumarate is a relatively mild electrophile, requiring millimolar concentrations to manifest substantial protein labeling (Fig. 1b). Treatment of proteomes with 2.5 mM fumarate caused near equivalent S-succination to that observed in *FH*−/− cell lines, consistent with previous reports suggesting fumarate accumulates to millimolar levels in HLRCC.^8^ Since antibodies can possess unanticipated cross reactivity and limited linear range, we devised an orthogonal chemical strategy to study fumarate’s covalent labeling using a clickable chemotype mimic, fumarate alkyne (FA-alkyne, Fig. 1c). FA-alkyne labeling occurred at slightly lower concentrations than fumarate-dependent S-succination in proteomes, potentially indicating the analogue’s heightened reactivity due to its lower LUMO energy (Fig. S1c). However, consistent with covalent labeling through a Michael addition mechanism, we observed time- and dose-dependent protein labeling upon incubation of lysates with FA-alkyne, but not the inert analogue succinate alkyne (Fig. S1d-e). FA-alkyne labeling was modestly competed by fumarate, but was completely abrogated by pre-incubation with MMF and DMF, again highlighting the attenuated reactivity of the oncometabolite relative to these molecules (Fig. S1f). Finally, we assessed fumarate’s reactivity in competitive labeling experiments using the established chemoproteomic reagent iodoacetamide alkyne (IA-alkyne),^26, 32^ and observed substantial blockade of cysteine labeling at low millimolar concentrations (Fig. 1d). Together, these results highlight the distinct reactivity of fumarate relative to DMF and MMF, and suggest this metabolite’s covalent reactivity may be most relevant in contexts such as HLRCC where it accumulates to millimolar levels.

### Global chemoproteomic profiling of fumarate-sensitive and FH-regulated cysteine residues

Next we set out to characterize novel targets of fumarate, employing a two-pronged chemoproteomic approach. First, we evaluated cysteine reactivity changes resulting from direct addition of exogenous fumarate to proteomes of an untransformed kidney cell line (HEK-293). These studies are designed to unambiguously define proteomic cysteine residues capable of directly reacting with fumarate, which we term “fumarate-sensitive” cysteines. Second, we applied chemoproteomics to map differential cysteine reactivity in FH-deficient (*FH*−/−) and FH-rescue (*FH+/+)* HLRCC cell lines, which we term “*FH*-regulated” cysteines. This latter comparison is mechanism-agnostic, identifying *FH*-regulated cysteine reactivity changes caused by direct S-succination, as well as alternative stimuli (such as oxidation or altered expression levels), and has potential to highlight biology specific to HLRCC cells.

To identify fumarate-sensitive and *FH*-regulated cysteines we applied IA-alkyne and an LC-MS/MS platform derived from isoTOP-ABPP (Fig. 2a-c).^32^ Briefly, paired samples consisting of either fumarate-treated and untreated proteomes, or *FH*−/− and *FH+/+* rescue HLRCC cells, were treated with IA-alkyne and conjugated to isotopically distinguishable azide biotin tags using click chemistry.

**Figure 2.**
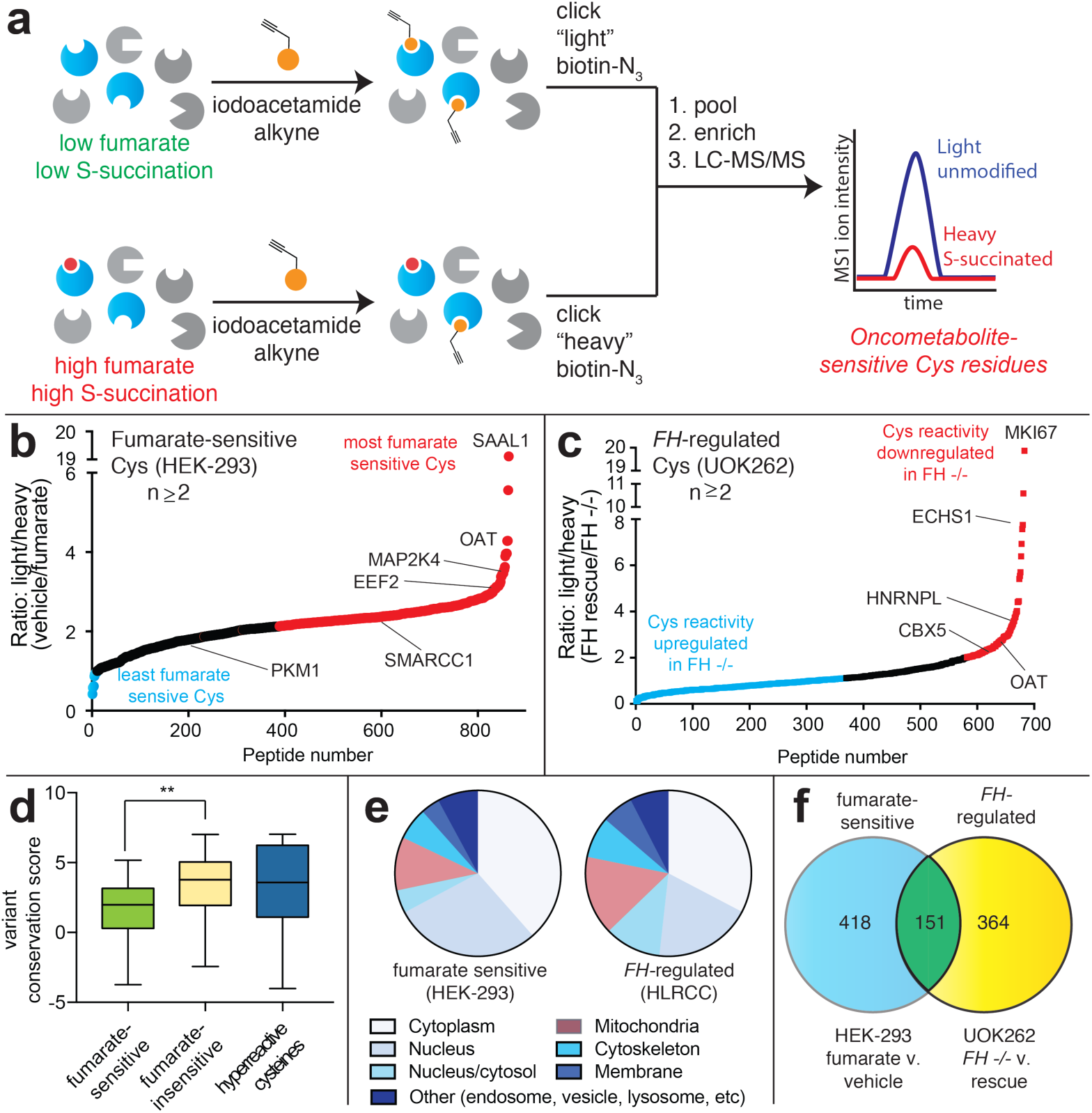
Global chemoproteomic profiling of fumarate-sensitive and *FH*-regulated cysteine residues. (a) Applying a competitive cysteine-profiling platform to study the oncometabolite fumarate. Experiments comparing untreated proteomes to those treated with exogenous fumarate define “fumarate-sensitive” Cys residues. Experiments comparing proteomes of *FH*−/− (UOK262) and *FH+/+* (UOK262WT) define “*FH*-regulated” Cys residues. (b) Fumarate-sensitive Cys residues identified in HEK-293 cells (n ≥2, SD ≤25%). (c) *FH*-regulated Cys residues identified in UOK262 cells (n ≥2, SD≤25%). (d) Conservation of fumarate-sensitive, fumarate-insensitive, and hyperreactive cysteine residues. (e) Subcellular localization of fumarate-sensitive and *FH*-regulated Cys residues. (f) Overlap of fumarate-sensitive Cys residues (R ≥2, n ≥2) with *FH*-regulated Cys residues identified in at least one experiment (R ≥1.2, n≥1).

Paired samples were pooled, enriched over streptavidin, subjected to on-bead tryptic digest, and IA-alkyne labeled peptides were released by dithionite cleavage of an azobenzene linker. LC-MS/MS was used to identify Cys-containing peptides, and the relative intensity ratio (ratio, R) of light/heavy isotopic pairs in the MS1 spectra was used as quantitative readout of Cys-labeling stoichiometry (Fig. 2a). In competitive fumarate labeling experiments, a L/H R value of ∼1 indicates that a cysteine was unaffected by fumarate, whereas a L/H R value of 2 indicates ∼50% modification (based on the formula “modification stoichiometry (%) = [1-(1/R)]*100%”; Fig. 2b). Analogously, in comparative analyses of HLRCC cells, a positive L/H R value indicates a cysteine’s reactivity (or abundance) is reduced by *FH* mutation (Fig. 2c).

Focusing first on fumarate-sensitive cysteines, we applied IA-alkyne to profile reactivity changes caused by treatment of proteomes with 1 mM fumarate for 15 hours (Fig. 2b, Table S1). This concentration was chosen to provide covalent labeling in the physiological range of S-succination but not saturate individual sites, which we hypothesized might limit the dynamic range of reactivity observed. We performed four independent replicate experiments, and applied additional reproducibility metrics to specify a subset of high confidence fumarate-sensitive cysteines (identified in ≥2 datasets, R standard deviation ≤25%). Using these criteria, we identified 854 cysteines out of >2300 quantified residues in human embryonic kidney proteomes which we characterized as fumarate-sensitive (Table S1). Of these cysteine residues, 569 were classified as highly sensitive (R≥2) and 285 as moderately sensitive (R = >1, <2), numbers which drop to 317 and 153, respectively, when requiring detection in ≥3 datasets. Identified among these hits were 25 known targets including *GAPDH*, which was found to be only moderately sensitive in our analysis, as well as *KEAP1* and *ACO2*, which did not meet the criteria for high confidence targets due to large standard deviations (Table S3). Comparing fumarate-sensitive cysteine residues to those identified in a recent chemoproteomic study of DMF,^30^ we find only a small fraction of residues with R≥ 2 (5.5%) overlap, once again highlighting the distinct reactivity of these molecules. An analysis of the evolutionary conservation of i) fumarate-sensitive cysteines, ii) fumarate-insensitive cysteines, and iii) hyperreactive cysteines^26^ revealed the former to be the least well-conserved (Fig. 2d). This is consistent with the hypothesis that fumarate may act as a covalent metabolite only upon hyperaccumulation, which would limit its reactivity from exerting strong evolutionary pressure. Along similar lines, the majority of fumarate-sensitive sites were identified on proteins found outside the mitochondria, where its concentration is highest (Fig. 2e). While in part this likely reflects the inherent challenge of analysis of the mitochondrial proteome,^33^ these data suggest mitochondrial enzymes may have evolved at an early stage to restrict reactivity with endogenous TCA cycle intermediates.

In order to complement our fumarate-sensitivity data, we next evaluated *FH*-regulated changes in cysteine reactivity (Fig. 2c). For these studies we applied IA-alkyne to comparatively profile an immortalized cell line derived from an HLRCC patient metastasis (UOK262 *FH*−/−) and a rescue cell line, in which the *FH* gene has been re-introduced through lentiviral transduction (UOK262WT, *FH+/+* rescue).^34^ Rescue cell lines exhibit reduced S-succination upon immunoblot analysis relative to *FH*−/− cells, consistent with their reduced fumarate levels and a measurable decrease in cysteine occupancy (Fig. S1a). Performing three independent replicate measurements of cysteine reactivity in this system led to the quantification of 1170 cysteine residues. Of *FH*-regulated cysteines whose reactivity changed ≥2-fold, 219 were upregulated and 112 were downregulated (Table S2). These numbers are reduced to 130 and 68 when requiring detection in multiple replicates, reflective of the overall increased noise observed in the endogenous system (Fig. 2c, Table S2).^35^ Comparison of our datasets revealed 151 cysteines classified as both *FH*-regulated and highly fumarate-sensitive (Fig. 2f). The remaining non-overlapping targets may result from differences in protein expression, oxidative modifications, and LC-MS/MS sampling between the two datasets. Comparing fumarate-sensitive and *FH*-regulated cysteines, we found the latter detected a higher percentage of mitochondrial proteins (Fig. 2e). This is consistent with the mitochondrial production of fumarate, and suggests reactivity-based profiling of *FH*−/− and *FH+/+* rescue HLRCC cells may sample distinct reactivity changes caused by oncometabolite compartmentalization. Interestingly, we found that individual proteins which incorporate multiple *FH*-regulated cysteines often exhibited unidirectional changes in their reactivity (Fig. S2a, Table S2), a profile that has previously been interpreted as indicative of global changes in protein abundance.^35^ However, we observed similar unidirectional cysteine reactivity changes in many proteins treated with exogenous fumarate (Fig. S2b, Table S1), suggesting this may be a genuine feature of covalent labeling by this mild electrophile.

To bolster our analysis, we further sought to identify proteins whose fumarate reactivity may be masked by altered protein abundance in *FH*−/− and *FH+/+* rescue HLRCC cells (Fig. 3). To examine this, we performed whole proteome (MudPIT) LC-MS/MS analyses of *FH*−/− and *FH+/+* rescue HLRCC cells and used this data to “correct” or normalize our reactivity measurements (Fig. S2c). Focusing on *FH*-regulated cysteines identified in ≥2 experiments, we obtained robust protein abundance data (≥10 spectral counts) for 53% of these parent proteins (Table S2). Using this data to correct our calculated reactivity for protein abundance led to modestly revised reactivity for the vast majority of residues analyzed, with 367/424 (86%) showing a less than 2-fold change. Of those that were altered, correction for abundance increased the calculated cysteine reactivity of 7 proteins by ≥2-fold, and decreased the calculated cysteine reactivity of 50 proteins by ≥2-fold (Table S2, Fig. S2d; an exemplary subset is illustrated in Fig. 3a). To validate our LC-MS/MS identifications, we assessed a subset of targets for fumarate-competitive labeling using the clickable chemotype mimic FA-alkyne (Fig. 3b). We analyzed proteins harboring fumarate-sensitive cysteines (EEF2, SMARCC1, MAP2K4), *FH*-regulated cysteines (OAT, HNRNPL), as well as abundance corrected *FH*-regulated cysteines (CBX5, Fig. S3d). Accordingly, lysates were incubated with fumarate, labeled with FA-alkyne, and subjected to click chemistry, enrichment, and Western blot (Fig. 3b). Consistent with LC-MS/MS data, capture of OAT, HNRNPL, EEF2, SMARCC1, and CBX5 exhibited modest to strong competition by fumarate treatment (Fig. 3c). In contrast, the non-target PKM1 showed no such competition. Interestingly, capture of MAP2K4, which was identified as harboring a fumarate-sensitive cysteine in our in vitro dataset, was also not sensitive to fumarate. This may indicate imperfect mimicry of fumarate’s labeling chemistry by FA-alkyne, the presence of multiple FA-alkyne reactive cysteine residues, or, alternatively, a false positive in our LC-MS/MS data. Overall, these studies demonstrate a strategy for comparing cysteine reactivity profiles in clonally distinct cell lines, and provide an initial glimpse into the sites and stoichiometry of the fumarate-reactive proteome.

**Figure 3.**
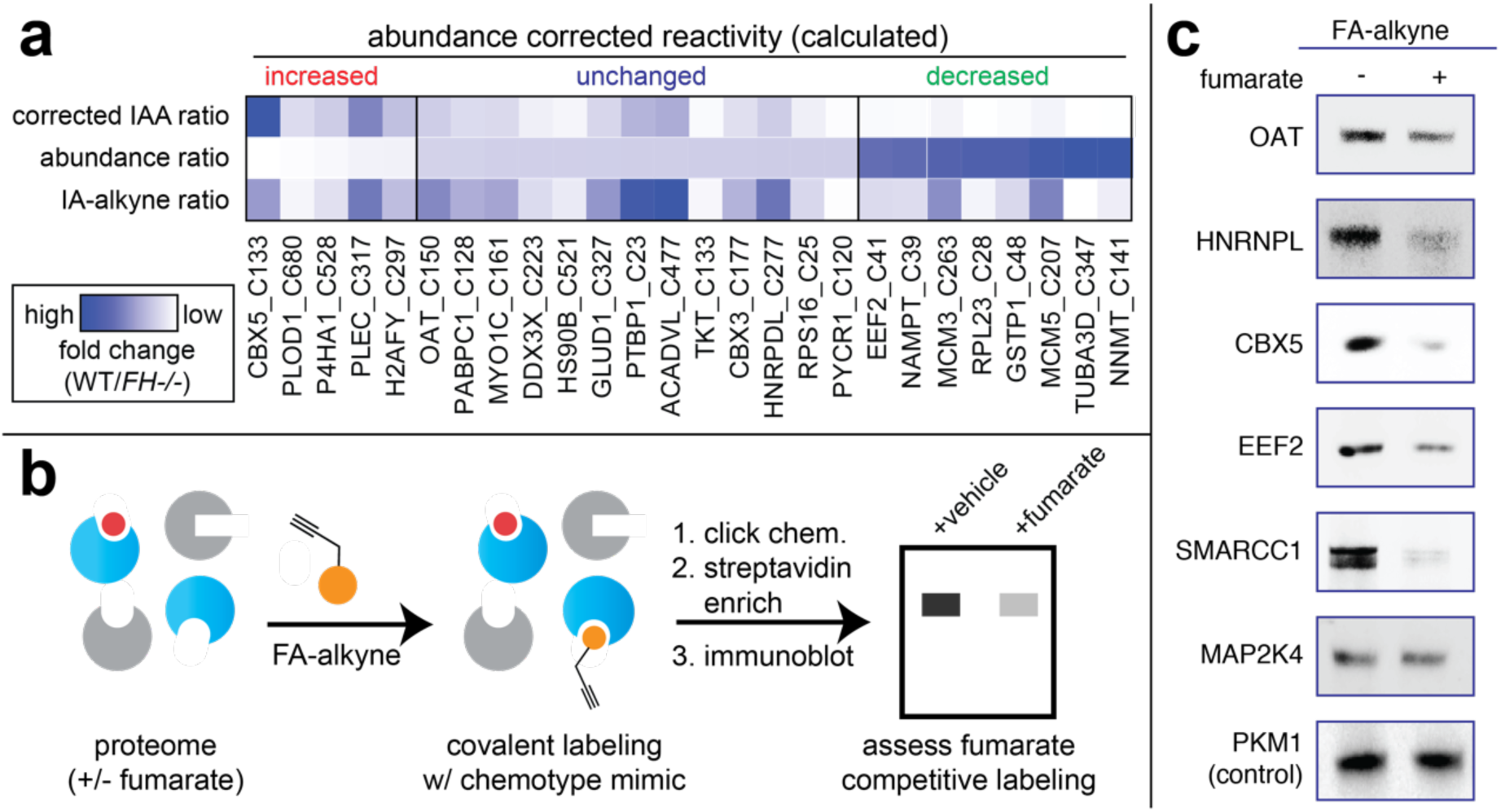
Analyzing the reactivity and abundance of *FH*-regulated cysteines. (a) Heat map illustrating strategy for correcting Cys reactivity ratios measured in UOK262 (FH-deficient) and UOK262WT (FH-rescue) cells using whole proteome MudPIT LC-MS/MS data. Adjusting for protein abundance can lead to an increase in calculated reactivity (left protein subset, red), an insignificant change (middle protein subset, blue), or a decrease in calculated reactivity (right protein subset, green). (b) Validating fumarate-sensitive and *FH*-regulated Cys residues using the clickable chemotype mimic FA-alkyne. (c) FA-alkyne capture of proteins that contain fumarate-sensitive or *FH*-regulated Cys residues is competed by fumarate (3 h pre-incubation with 1 mM fumarate; then 15 h treatment with 100 μM FA-alkyne).

### Molecular and structural determinants of covalent fumarate-cysteine interactions

Next we sought to utilize our knowledge of fumarate-sensitive cysteines to better understand the structural determinants of oncometabolite reactivity. As an initial step, we assessed the highest stoichiometry fumarate-sensitive cysteines (which represent candidate direct targets of fumarate) for the presence of linear motifs in their flanking sequences using pLogo (Fig. 4a, Table S4).^36^ Interestingly, fumarate-sensitive cysteines showed an enrichment of acidic residues such as glutamate (E) and aspartate (D) in their flanking regions. This was unexpected, as nucleophilic cysteines typically are surrounded by proximal basic residues such as lysine (K) and arginine (R), which can serve as hydrogen bond donors and help stabilize the developing negative charge of the thiolate. The atypical nature of this fumarate-sensitivity motif was further supported by pLogo analysis of fumarate-insensitive and hyperreactive cysteine residues,^26^ each of which demonstrated the expected enrichment of basic, rather than acidic, residues in their flanking motif (Fig. 4b, Fig. S3a). Fumarate’s cysteine reactivity motif was also distinct from that of DMF and HNE (Fig. S3a),^30, 37^ with only MMF showing similar enrichment of flanking acidic residues (Fig. S3a). Considering the hypothesis that fumarate-sensitive cysteines may possess a unique local sequence environment, we next asked how fumarate-sensitivity correlated with overall cysteine reactivity. For this analysis we overlaid upon our fumarate-sensitive cysteines a previously constructed map of proteome-wide cysteine reactivity, constructed from concentration-dependent analysis of IA-alkyne labeling.^26^ In contrast to stimuli such as DMF^30^ or GSNO,^32^ which target cysteine residues across the reactivity spectrum, fumarate-sensitive cysteines are strikingly anti-correlated with reactivity (Fig. 4c, Fig. S3b-c). *FH*-regulated cysteines identified in HLRCC cells exhibit a similar anti-correlation, suggesting this is not an artifact of the competitive labeling experiment (Fig. 4d). Furthermore, in *GSTO1*, a model gene which harbors a known nucleophilic cysteine residue, fumarate was found to preferentially impact a distal cysteine while leaving the active site residue unmodified (Fig. 4e). Similar observations were made for the hyperreactive cysteine of *NIT2* during analysis of *FH*-regulated cysteines (Fig. S3b). These analyses define a unique local environment for covalent oncometabolite labeling.

**Figure 4.**
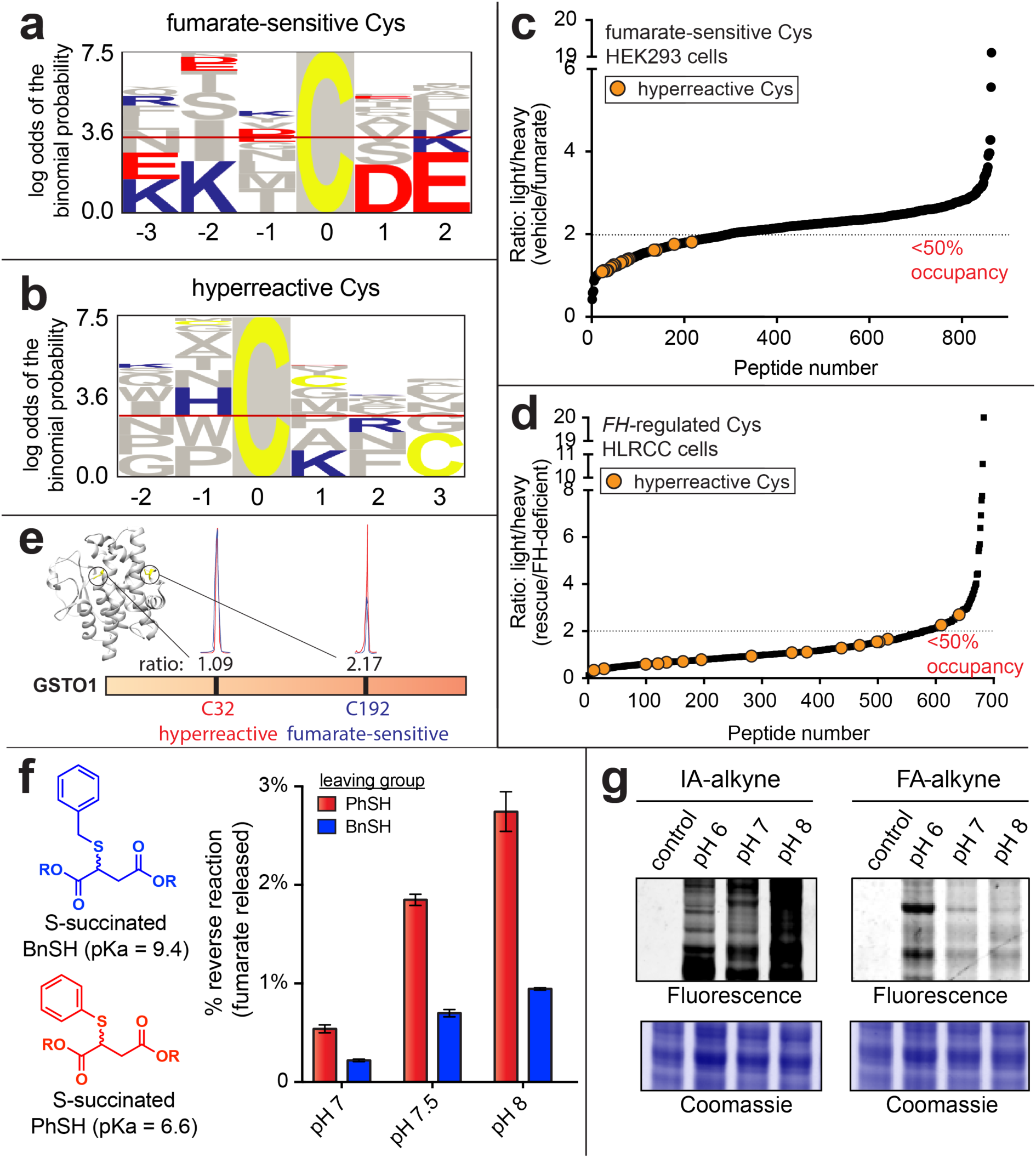
Establishing the molecular determinants of covalent fumarate-protein interactions. (a) Motif analysis of fumarate-sensitive Cys residues reveals an enrichment in flanking carboxylates. (b) Motif analysis of hyperreactive Cys residues identified in Weerapana et al.^26^ (c) Fumarate-sensitive Cys residues are anti-correlated with Cys reactivity. (d) *FH*-regulated Cys residues are anti-correlated with Cys-reactivity. (e) Fumarate preferentially targets a less nucleophilic, non-active site Cys in GSTO1. (f) The reversibility of S-succinated model thiols is dependent on leaving group pKa, but does not proceed to an appreciable extent. S-succinated thiols (1 mM) were incubated in 100 mM Tris buffers at 37 °C for 24 h, prior to quantification of DMF release by fluorescence assay. (g) In contrast to neutral electrophiles such as IA-alkyne, FA-alkyne exhibits a paradoxical increased reactivity at lower (more acidic) pH. Probe treatments were carried out as follows, prior to the click chemistry-left: IA-alkyne (1h; 100 μM); right: FA-alkyne (15 h, 1 mM).

In order to better understand the mechanistic basis for these observations, we explored two hypotheses. First, we assessed the pKa-dependent reversibility of cysteine S-succination. Our rationale was that, if low pKa hyperreactive cysteines act as better leaving groups in retro-Michael reactions, then irreversible reaction with IA-alkyne may obscure their reversible reaction with fumarate, leading to the observed anti-correlation. To test this hypothesis, we synthesized S-succinated thiols derived from mercaptans of disparate acidities (Fig. 4f). Next, we assessed the reversibility of these model substrates using a recently developed fluorescence assay to monitor fumarate release (Fig. S3e-f).^38^ This experiment indicated that while S-succinated thiols did indeed exhibit pKa-dependent reversibility, the extent of fumarate release was minor, with only 2-4% reversal observed over 24 hours (Fig. 4f, Fig. S3g). This is consistent with previous studies^39^ as well as the structure of S-succinated cysteine, which resembles ring-opened maleimides engineered for stable covalent labeling of protein cysteine residues.^40^ These studies suggest that while reversible S-succination is possible, it makes a negligible or minor contribution to fumarate’s unique covalent labeling profile.

Next, we considered two ways in which electrostatics may alter fumarate reactivity. First, the anionic nature of fumarate may favor its association with protein surfaces, which are enriched in charged residues, compared to more hydrophobic hyperreactive cysteine-containing active sites. Indeed, previous studies have observed unique sites of protein labeling for negatively charged electrophiles relative to their neutral analogues.^41-43^ Second, we hypothesized that protonated hydrogen fumarate may function as the active electrophile in S-succination reactions. Protonation of fumarate would increase fumarate’s reactivity by lowering its LUMO energy (Fig. S3h), and furthermore limit repulsive electrostatic interactions with negatively charged thiolates and active-site proximal carboxylates. Speaking to the feasibility of such a mechanism, the viability of hydrogen fumarate as a reactive species in aqueous buffer has precedence in previous studies of MMF reactivity.^31^ To explore the potential relevance of these phenomena, we tested the influence of pH on fumarate reactivity. The neutrally charged electrophile IA-alkyne exhibits increased protein labeling at higher pH, presumably due to higher thiolate concentrations (Fig. 4g). In contrast, fumarate and FA-alkyne cause increased protein labeling at lower pH, conditions which favor decreased thiolate and increased hydrogen fumarate concentrations (Fig. 4g, Fig. S3i). Addition of high salt to S-succination reactions, which would be expected to decrease fumarate’s pKa and reduce ionic interactions, led to overall reduced labeling of proteins by fumarate (Fig. S3j). In addition to revealing a paradoxical influence of pH on fumarate reactivity, these studies highlight hydrogen fumarate as a novel molecular entity potentially responsible for covalent protein modification in HLRCC.

### Fumarate-sensitive and FH-regulated cysteines lie in several kidney cancer pathways

To identify novel biology affected by covalent oncometabolism in HLRCC, we next performed pathway analysis of *FH*-regulated and fumarate-sensitive cysteines. In order to focus our analysis on succination events likely to have functional effects on protein activity we employed the informatics tool Mutation Assessor, which predicts functional mutations on the basis of sequence conservation.^44^ C to E mutations of fumarate-sensitive cysteine residues were used to mimic the negative charge gained upon covalent S-succination. Analysis of highly fumarate-sensitive cysteines (R≥2, 50% stoichiometry) identified by competitive chemoproteomics in all four replicates highlighted 58 proteins whose modification was expected to have a high or moderate impact on protein function (Fig. 5a, Table S4). Applying a similar workflow to *FH*-regulated cysteines whose reactivity was downregulated in HLRCC cells (R≥2, 50% stoichiometry) led to the identification of an additional 69 proteins. Gene ontology analysis found that these candidate S-succinated proteins clustered in pathways related to mitochondria, metabolism, RNA processing, and transcriptional regulation, many of which play known roles in kidney cancer pathogenesis (Fig. 5b, Fig. S4a). This suggests several novel cellular pathways which may be deleteriously impacted by fumarate’s covalent reactivity in HLRCC. To prioritize targets for mechanistic follow-up we used two criteria. First, we focused on genes for whom multiple patients harboring loss-of-function genetic lesions (mutations or deletions) had been identified in kidney cancer sequencing experiments (Fig. 5c),^45^ positing these targets may be relevant to renal cell tumorigenesis. Second, we prioritized genes for which structural data indicated the candidate fumarate-sensitive cysteine mapped to a predicted cofactor or protein-protein interaction site (Fig. S4b-i), which we hypothesized would facilitate mechanistic analyses. Employing these criteria, *SMARCC1* emerged as 1) the fumarate-sensitive gene most commonly lost in renal cell carcinoma patients (Fig. 5c), and 2) harboring a site of fumarate-sensitivity near a protein-protein interaction interface (Fig. S4b, Fig. 5d-e), This led us to further focus on SMARCC1 as a case study to understand the functional consequences of fumarate reactivity.

**Figure 5.**
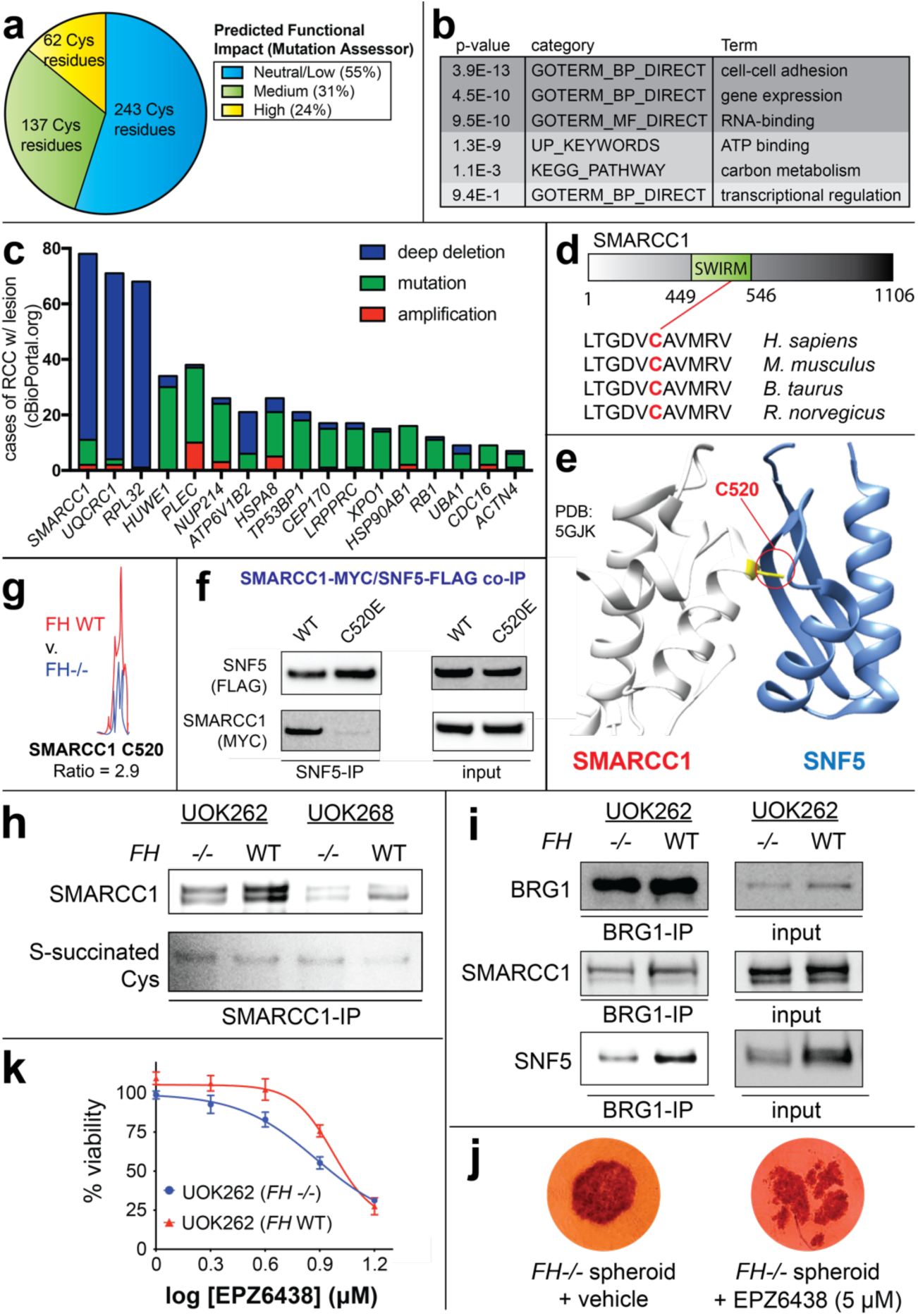
Functional analyses of fumarate-sensitive Cys residues. (a) Percentage of fumarate-sensitive and *FH*-regulated Cys residues predicted to be functional using the informatics tool Mutation Assessor. (b) Gene ontology analysis of fumarate-sensitive and *FH*-regulated Cys residues. (c) Correlation between genes containing fumarate-sensitive Cys residues and genetic alterations in kidney cancer (renal cell carcinoma, RCC). (d) SMARCC1 C520 lies in the SWIRM domain and is conserved in higher organisms. (e) SMARCC1 C520 lies at the SNF5 subunit interface. (f) SMARCC1 C520E mutation limits co-immunoprecipitation with SNF5 in HEK-293 cells co-overexpressing FLAG-tagged SNF5 with Myc-tagged SMARCC1 (WT or C520E mutant). (g) SMARCC1 C520 undergoes FH-dependent changes in occupancy in HLRCC cell lines despite identical expression levels [Figure S5]. (h) SMARCC1 S-succination can be detected in FH-deficient and FH WT HLRCC cell lines post-immunoprecipitation of endogenous SMARCC1. (i) SNF5 demonstrates decreased co-immunoprecipitation and decreased levels in *FH*−/− HLRCC cells. Left: Results from SWI/SNF complex co-immunoprecipitation with BRG1 antibody. Right: Endogenous levels of SWI/SNF complex members in HLRCC cells. (j) EZH2 inhibitors are toxic to HLRCC spheroids. UOK262 *FH* −/− spheroids were treated with vehicle or EPZ6438 (14 days; 5 μM). Figure shown here is representative of 6 replicates. (k) EZH2 inhibitors exhibit modest selectivity for FH-deficient HLRCC cells. UOK262 *FH* −/− or *FH* WT spheroids were treated with EPZ6438 (21 days; 1, 2, 4, 8, 16 μM) and % viability plotted relative to the vehicle treated spheroids.

### Functional analysis of a fumarate-reactive cysteine in SWI/SNF complex

*SMARCC1* is a core member of the SWI-SNF chromatin remodeling complex, a known tumor suppressor in many cancers.^46-47^ *SMARCC1* is commonly deleted in clear cell renal cell carcinoma (ccRCC) due to its position on the short arm of chromosome 3, which lies adjacent to the *VHL* tumor suppressor. Of note, *SMARCC1* does not exhibit coordinate mutation and deletion in *VHL*-deficient ccRCC, suggesting an intact genomic copy of *SMARCC1* is required for cell growth.^48^ Among the nine cysteine residues on SMARCC1, Cys520 (C520) was exclusively identified as fumarate-sensitive by competitive chemoproteomic experiments. Cys520 lies in SMARCC1’s SWIRM domain, the most common site of SMARCC1 somatic mutation in cancer. Studies of SMARCC1’s mouse ortholog (Srg3) have found the SWIRM domain regulates the stability of SNF5, a tumor suppressive subunit of SWI/SNF, via direct protein-protein interactions.^49^ A recent crystal structure revealed C520 lies within a solvent-exposed helix residing directly at the SMARCC1-SNF5 interface, suggesting its modification may obstruct this protein-protein interaction (Fig. 5e).^50^ To test this hypothesis, we performed co-immunoprecipitation experiments in HEK-293 cells transfected with plasmids encoding FLAG-tagged SNF5 and either wild-type SMARCC1, or a C520E mutant. Mutation of C520 was found to completely abrogate the ability of SNF5 to capture SMARCC1 (Fig. 5f, S5a). Similarly, while ectopic expression of wild type SMARCC1 stabilized SNF5, the C520E mutation had less of an effect (Fig. S5b).^51^ Additionally, we found that treatment of cells co-overexpressing SMARCC1/SNF5 with cell-permeable ethyl fumarate also reduced SNF5 stability (Fig. S5c).

Analysis of HLRCC cells revealed greater labeling of SMARCC1 C520 by IA-alkyne in *FH+/+* rescue as compared to *FH*−/− cells, consistent with covalent modification of this residue by endogenous fumarate (Fig. 5g). We also identified evidence for modification of the homologous cysteine in SMARCC2, whose SWIRM domain is nearly identical to SMARCC1’s (Fig. S5d-e). However, direct detection of the S-succinated C520 proved more challenging. MudPIT with PTM analysis of *FH*−/− cell lines validated S-succination of several novel proteins identified in our two datasets (GCLM, PCBP1, TCP1, Table S5), but not SMARCC1. S-succination blots of SMARCC1 immunoprecipitated from *FH*−/− and *FH+/+* rescue HLRCC cells were characterized by high background, with slightly increased signal in FH-deficient cells (Fig. 5h). To further explore this phenomenon, we next examined HLRCC cells for evidence of a disrupted SMARCC1-SNF5 interaction. Co-immunoprecipitation of SWI/SNF complex indicated a modest decrease in SMARCC1’s interaction with SNF5 in *FH*−/− relative to *FH+/+* rescue cells (Fig. 5i). In line with decreased interaction, SNF5 protein, but not transcript levels, are also lower in these cells (Fig. 5i, Fig. S5f-g). While the interaction between SMARCC1 and SNF5 is weakened, it is not fully disrupted, as glycerol gradient fractionation indicated that the core SWI/SNF complex remains intact (Fig. S5h). Finally, we observe that the EZH2 inhibitor EPZ6438 (tazematostat) manifests a FH-dependent toxicity in HLRCC spheroids (Fig. 5 j-k). Of note, decreased SNF5 levels have previously been shown to sensitize tumors to EZH2 inhibition,^52^ and the observed effect is consistent with a measurable, but minor, effect of fumarate on SNF5 function. These studies demonstrate a novel cysteine-dependent protein-protein interaction in the SWI-SNF complex that may be modulated by oncometabolite accumulation.

### Comparative chemoproteomics reveals ligandable cysteines upregulated in HLRCC

Covalent modifications driven by metabolite reactivity are largely expected to exert deleterious effects on protein function.^53^ However, as a final experiment we wondered whether chemoproteomic analyses may also be capable of identifying pathways positively influenced by *FH* mutation. To explore this idea we re-analyzed our chemoproteomic data from *FH*−/− and *FH*+/+ rescue HLRCC cell lines, particularly focusing on *FH*-regulated cysteines with R values <1 (blue region, Fig. 2c). Cysteines with these values are almost exclusively found in our comparative analysis of HLRCC cells, and are expected to originate from proteins whose abundance or activity is increased by FH-deficiency. We applied gene set enrichment analysis (GSEA)^54^ to these datasets, and found that FH-deficient cells enriched reactive cysteines belonging to proteins activated by the transcription factors HIF-1*α* and NRF2 (Fig. 6a). This was notable, as each of these pathways have previously been shown to be overactive in HLRCC.^7, 17, 19^ This suggests chemoproteomic analyses may provide a useful complement to traditional methods such as gene expression profiling for the discovery of novel cancer pathways. However, an additional advantage of these approaches is that they also have the potential to identify leads for covalent ligand development, as powerfully demonstrated in a recent analysis of NRF2 mutant cancers.^35^ To facilitate such efforts, we cross referenced cysteines activated by *FH* mutation with a recently disclosed covalent fragment library whose proteome-wide targets were characterized.^28^ This analysis identified ligandable cysteines in many proteins lying within pathways upregulated upon *FH* loss, including glycolysis, hypoxia, and reactive-oxygen stress (Fig. 6b). These studies highlight a strategy for mining chemoproteomic data to identify novel targets and lead fragments for pathway disruption in HLRCC.

**Figure 6.**
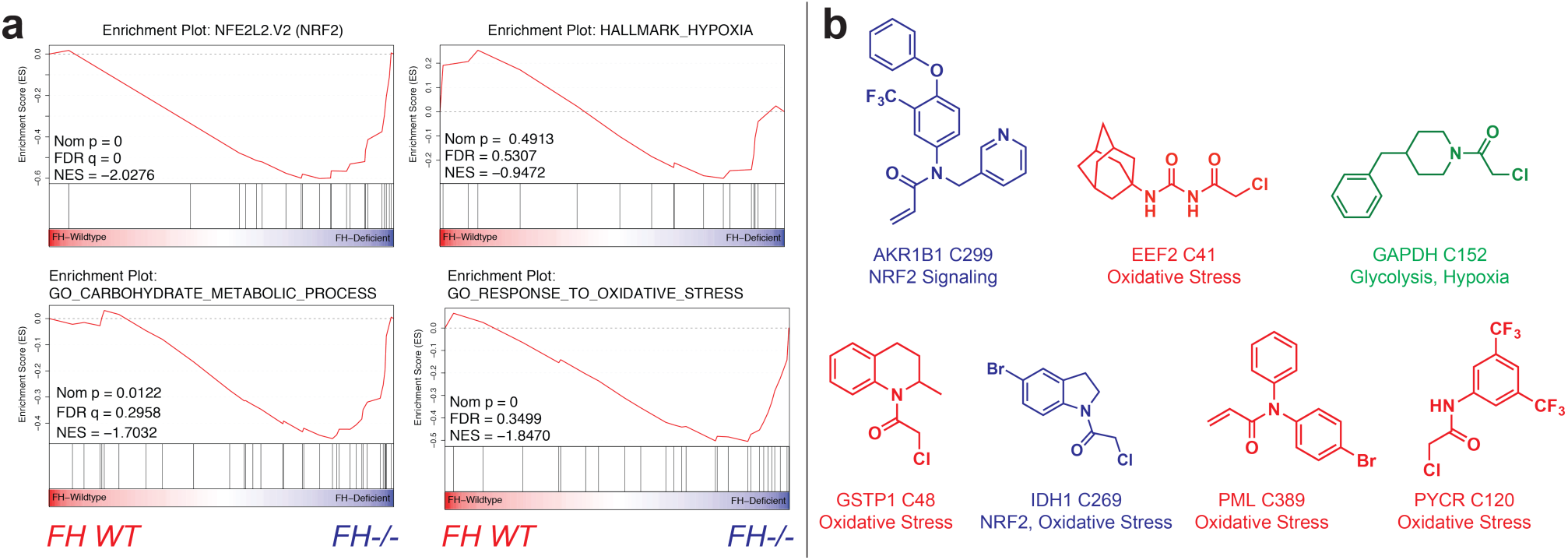
(a) Gene set enrichment analysis (GSEA) of *FH-*regulated Cys residues highlights pathways whose activity is functionally upregulated in HLRCC cells. (b) Fragments identified by Backus et al. targeting Cys residues whose reactivity is enhanced in FH-deficient HLRCC cells.

## Discussion

The discovery of hereditary cancers driven by TCA cycle mutations provided some of the first evidence that metabolites themselves could fuel tumorigenic signaling.^1-3^ Recent data indicates that “oncometabolic” signaling is not unique to these contexts, but may instead represent a broader aspect of malignancy.^9, 18^ While this provides another powerful example of how studying genetic disorders can illuminate general cancer mechanisms,^55^ understanding precisely how oncometabolites drive malignant transformation and developing diagnostics to track this process remains a critical goal. Here we have applied chemoproteomics to define a novel complement of protein cysteines sensitive to the oncometabolite fumarate and *FH* mutation in the genetic cancer syndrome HLRCC. Competitive labeling using IA-alkyne facilitates the discovery of fumarate-sensitive cysteines while circumventing the necessity for antibody-generation or direct detection of poorly ionizable, negatively charged S-succinated peptides. A distinct feature of our strategy was the parallel identification of both “fumarate-sensitive” targets via direct competitive labeling, as well as endogenous “*FH*-regulated” targets via comparative profiling of *FH*−/− or *FH+/+* rescue HLRCC cell lines. To our knowledge, this represents the first application of reactivity-based protein profiling to study an endogenous covalent metabolite,^37, 56^ as well as the mutational context that causes its production.^35^ In the future, we anticipate the analyses presented here should provide a useful model for the study of mutational and non-mutational stimuli which cause production of other endogenous electrophiles, such as lipid electrophiles,^37^ itaconate,^57-58^ and acyl-CoAs.^59-60^

Fumarate’s mild reactivity is distinct from that of endogenous species such as hydrogen peroxide and hydroxynonenal whose cysteine reactivity has previously been profiled.^37, 56^ It is important to note that while substantial evidence points to an important mechanistic contribution of fumarate’s covalent reactivity to HLRCC pathogenesis, our chemoproteomic data has other important applications. For example, enzymes containing cysteines whose abundance or reactivity is increased upon *FH* loss may represent activities whose upregulation is necessary for HLRCC survival (Fig. 6). Alternatively, it is plausible that a subset of functional fumarate-sensitive cysteines do not facilitate tumorigenesis, but rather create collateral vulnerabilities that may be exploited for therapy.^61^ This is analogous to the manner in which BRCA1 mutation sensitizes breast cancer to PARP inhibition,^62^ and is a strategy that has already demonstrated utility in the targeting of oncometabolite-dependent cancers.^63-65^ Pathways enriched in functional fumarate-sensitive cysteines in our gene ontology analyses represent attractive candidates for exploration of this paradigm (Fig. 5a, Table S4). Finally, non-functional, but high stoichiometry targets of fumarate may provide stage-specific biomarkers of *FH* status in HLRCC,^20^ potentially detectable by site-specific antibody generation or targeted LC-MS/MS assays. In addition to their relevance to HLRCC, we anticipate these data will provide a rich resource for understanding fumarate’s covalent reactivity in other settings where this metabolite’s accumulation has been observed, including diabetes,^15, 66^ non-hereditary kidney cancer,^67-68^ neuroblastoma,^69^ colorectal cancer,^70-71^ and tumors of the adrenal gland.^68^

An unanticipated finding of our studies was the observation that fumarate-sensitive cysteines are enriched in flanking carboxylates and demonstrate an anti-correlation with cysteine reactivity. Of note, this relationship was not observed with other cysteine-reactive electrophiles. Furthermore, in contrast to iodoacetamide, fumarate (as well as its mono-carboxylate analogue FA-alkyne) showed increased protein reactivity at acidic pH. These observations recall a recent study of MMF, which upon incubation with GSH was found to form an equimolar mixture of regioisomeric adducts at the C2 and C3 positions, consistent with protonated hydrogen MMF acting as the electrophile.^31^ By analogy, the observed pH-dependence suggests that protonation of fumarate, rather than deprotonation of cysteine, is rate-limiting for fumarate reactivity. This theory, arising from our chemoproteomic data, suggests that hydrogen fumarate, rather than fumarate, is the signature covalent oncometabolite of HLRCC, and has several implications. First, since the concentration of hydrogen fumarate is extremely low in physiological buffer, it suggests that fumarate is likely to act as a covalent metabolite only under conditions which favor its hyperaccumulation as well as acidic pH. In this regard HLRCC tumors provide a near ideal environment for S-succination, as in addition to high fumarate caused by disruption of FH, they exhibit potent production of lactic acid due to increased glycolytic metabolism.^7, 34^ An additional, technical implication is that the repertoire of reactivity-based protein profiling methods,^26, 28^ as well as covalent cysteine-targeting ligands, may be expanded by incorporating electrostatics as a design element. Future studies will be required to test this hypothesis, as well as better understand the origins of fumarate’s distinct reactivity.

Fumarate-sensitive cysteines were found to reside in functional domains of proteins that mapped to a number of pathways relevant to kidney cancer, including mitochondrial metabolism, RNA processing, and gene expression. Mechanistic analyses of SMARCC1 discovered a fumarate-sensitive cysteine in the protein’s highly-conserved SWIRM domain that is critical for protein-protein interactions with the tumor suppressor SNF5. Furthermore, we found that HLRCC cells exhibit FH-dependent changes in SMARCC1 C520 occupancy, as well as evidence of modest SWI/SNF dysfunction, including reduced SMARCC1-SNF5 interaction, diminished SNF5 levels, and susceptibility to EZH2 inhibitors. In addition to defining a minor impact of fumarate on SWI/SNF in HLRCC, our findings also illustrate a larger concept, specifically the potential of covalent metabolites to regulate protein-protein interactions (PPIs) in the nucleus. Building on our analysis of fumarate’s unique labeling profile, is tempting to speculate that the polar labeling environment preferred by fumarate-sensitive cysteines may predispose the high stoichiometry display of S-succination towards solvent-exposed surfaces capable of disrupting biomolecular interactions. While relatively few examples of non-enzymatic reactive metabolite-dependent PPIs exist,^53^ the role of cysteine oxidation in regulating such interactions is well-precedented.^56, 72-73^ Our studies suggest further investigation of fumarate-dependent protein-protein and protein-nucleic acid interactions is warranted.

Finally, it is important to point out some limitations of our current approach, as well as future avenues that may help to address them. One important drawback of our competitive profiling method is that it does not directly identify S-succinated cysteines, but rather detects cysteines whose reactivity is altered by fumarate treatment or *FH* mutation. This leads to the caveat that the observed reactivity changes could be due to direct modification by fumarate, or alternative species such as lipid electrophiles or reactive oxygen species that are known to be produced as a consequence of *FH* mutation.^17, 74^ Therefore, an important future goal will be the development of improved methods for direct S-succination analysis, including immunoprecipitation-grade antibodies, and/or techniques analogous to the biotin-switch protocols used to investigate other cysteine modifications.^75^ In addition, our work illustrates the difficulty of differentiating between changes in reactivity and expression in *FH* mutant and wild-type cell lines. This issue was also encountered in a recent study of *NRF2* mutant and wild-type lung cancers^35^, which normalized cysteine reactivity to RNA-Seq gene expression, in contrast to the whole proteome MudPIT LC-MS/MS data used here. Integration of these two complementary approaches will likely provide a more complete picture of cysteine reactivity in future comparative profiling studies. Finally, a critical challenge illustrated by the current study arises from the sheer magnitude of candidate targets. Using a reproducibility metric of replicate detection, we identified >100 cysteines predicted to be functionally impacted by fumarate’s reactivity (Table S4), a number which rapidly increases when requiring detection in only a single dataset. While relatively small on a genomic scale, this number of targets far exceeds the bandwidth of most laboratories for mechanistic follow-up. Therefore, in the future we anticipate chemoproteomic studies of pleiotropic metabolites such as fumarate will benefit from marriage to other high-throughput methods, such as pooled CRISPR or siRNA screening approaches^76^, which may allow high-throughput validation of oncometabolite targets. The data presented here will provide an information-rich resource for such studies, and ultimately facilitate the definition of fumarate as a signaling molecule and biomarker in HLRCC and other pathological settings marked by oncometabolite accumulation.

## Supplemental Tables

**Table S1.** Fumarate-sensitive cysteines identified by competitive profiling HEK-293 cells treated and untreated with fumarate.

**Table S2.** *FH*-regulated cysteines identified by comparative profiling of *FH*−/− HLRCC cell line (UOK262) and a *FH+/+* rescue HLRCC cell line (UOK262WT).

**Table S3.** Compiled list of S-succinated cysteine residues previously characterized in the literature, and annotation with chemoproteomic data (if available).

**Table S4.** Sequences used for motif analysis, as well as results for analyses of conservation-based functional impact (FI), gene ontology (GO), and genomic lesions found in covalent fumarate targets in kidney cancer.

**Table S5.** Peptides identified as targets of S-succination in MudPIT LC-MS/MS analyses of HLRCC cell (UOK262 and UOK268) proteomes.

## Figure Supplements

**Figure S1.**
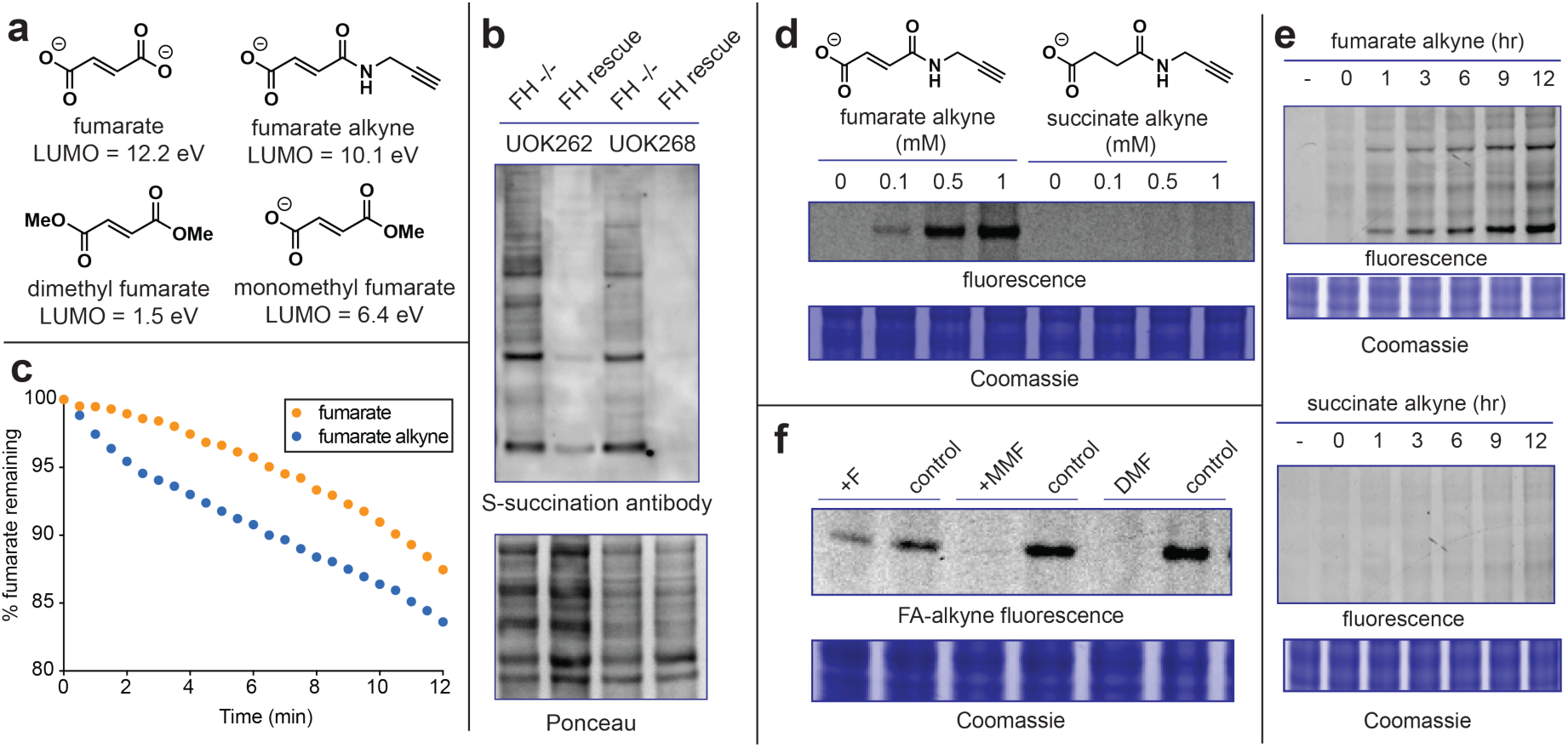
(a) Lowest unoccupied molecular orbital (LUMO) energies of fumarate analogues. (b) Cysteine S-succination in HLRCC cells is FH-dependent. (c) Relative reactivity of fumarate (1 mM) and fumarate alkyne (FA-alkyne, 1 mM) with a model thiol (2-mercaptoethanol, 10 mM) in PBS as measured by UV analysis of fumarate at 240 nm. (d) FA-alkyne but not succinate alkyne exhibits dose-dependent protein labeling (15 h treatment; 100, 500, 1000 μM). (e) FA-alkyne (1 mM; top panel) but not succinate alkyne (1 mM; bottom panel) exhibits time-dependent protein labeling (0, 1, 3, 6, 9, 12 h treatment). (f) Competitive FA-alkyne labeling reveals the reactivity of fumarate (F) is attenuated relative to the previously studied drug dimethyl fumarate (DMF), as well as its metabolite monomethyl fumarate (MMF), (3 h pre-incubation with 1 mM competitor; then 15 h treatment with 100 μM FA-alkyne).

**Figure S2.**
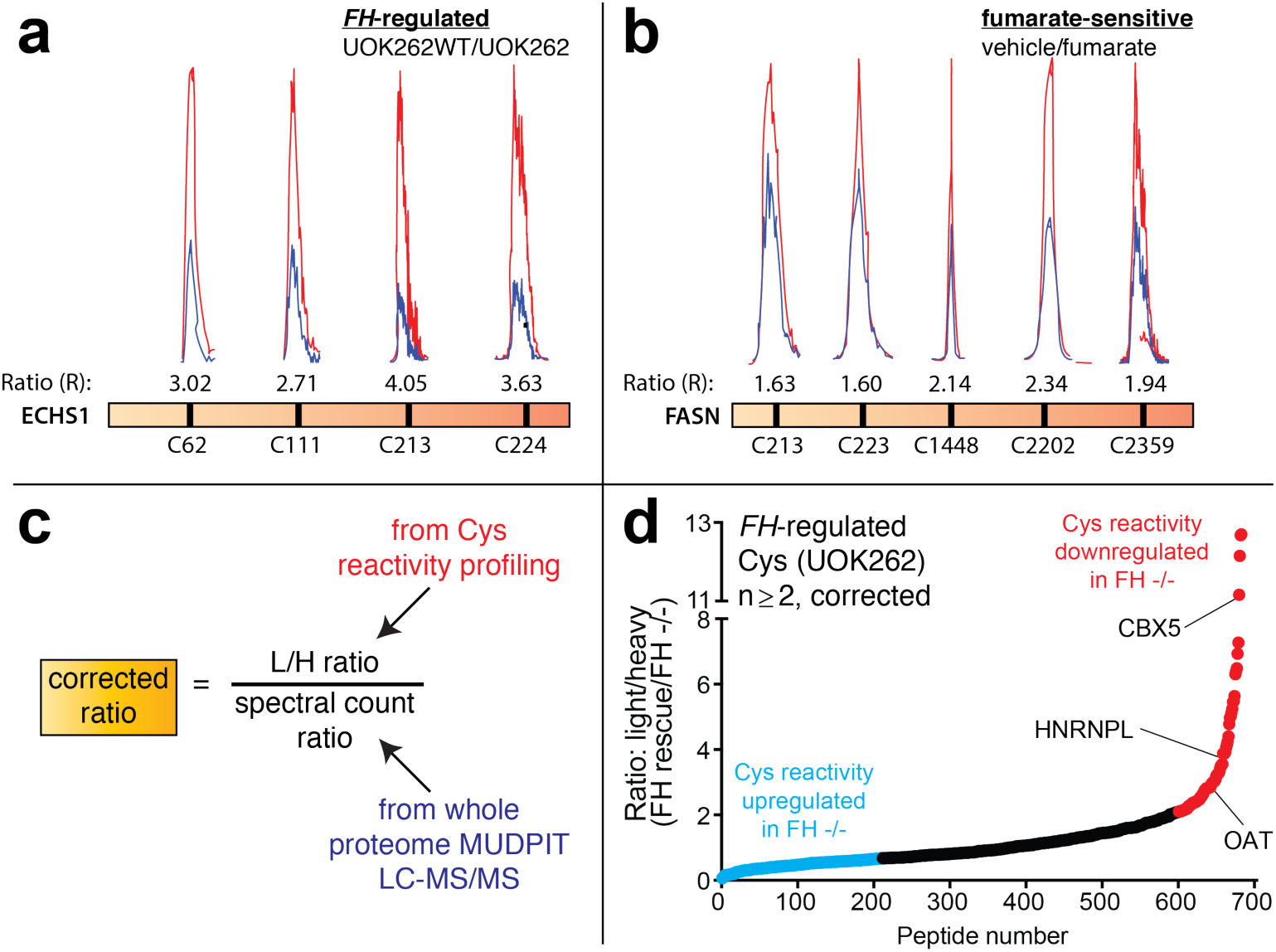
Examples of proteins with multiple Cys residues exhibiting unidirectional reactivity changes in (a) *FH*-regulated, and (b) fumarate-sensitive datasets. (c) Calculation for corrected Cys reactivity in *FH*-regulated datasets using whole proteome MudPIT data. (d) Plot of *FH*-regulated cysteine residues correcting for abundance changes observed by MudPIT.

**Figure S3.**
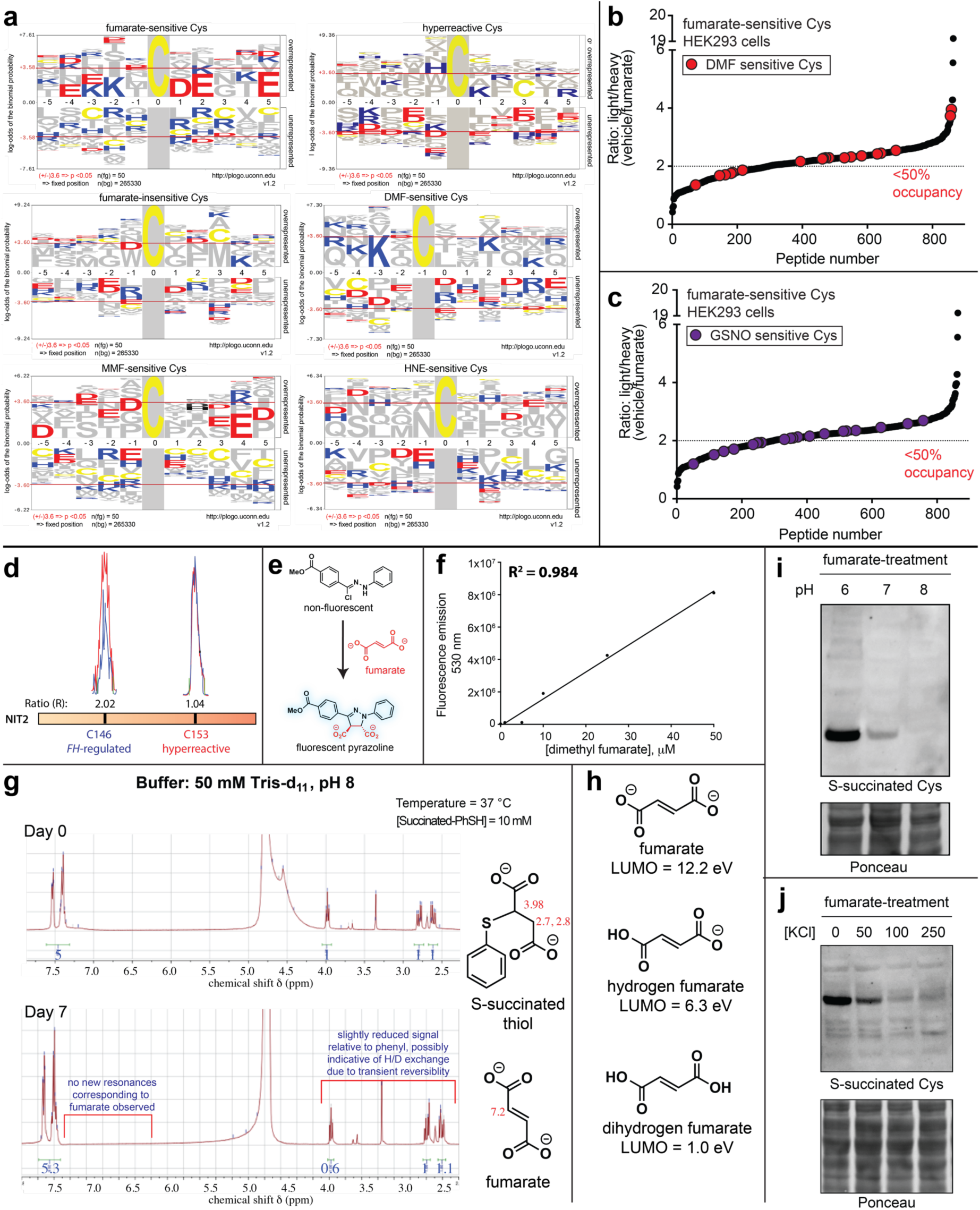
(a) Motif analysis for (left to right) fumarate-sensitive cysteines, hyperreactive cysteines (Weerapana et al.), fumarate-insensitive cysteines, DMF-sensitive cysteines, MMF-sensitive cysteines, and 4-hydroxynonenal (HNE)-sensitive cysteines. (b) and (c) DMF and GSNO target cysteines across the fumarate-sensitivity spectrum. (d) Fumarate preferentially targets a less nucleophilic, non-active site Cys in NIT2. (e) Fluorogenic reaction for fumarate detection. (f) Calibration curve for fluorescent detection of DMF release from model S-succinated substrates. (g) NMR analysis of reversible S-succination. ^1^H NMR spectra of model S-succinated thiols (10 mM final concentration, 50 μL of 100 mM stock in DMSO-*d_6_*) incubated in 50 mM TRIS-*d_11_* buffer (pH 8, adjusted using 1 M deuterium chloride, 450 μL) in D_2_O at 37 °C for 7 days. (h) LUMO energies of fumarate and protonated analogues. (i) Proteomic S-succination proceeds more efficiently at lower pH. (j) Proteomic S-succination by exogenous fumarate is antagonized by increasing ionic strength. For (i) and (j) HEK-293 proteomes were treated with fumarate (5 mM) for 15 h prior to western blotting.

**Figure S4.**
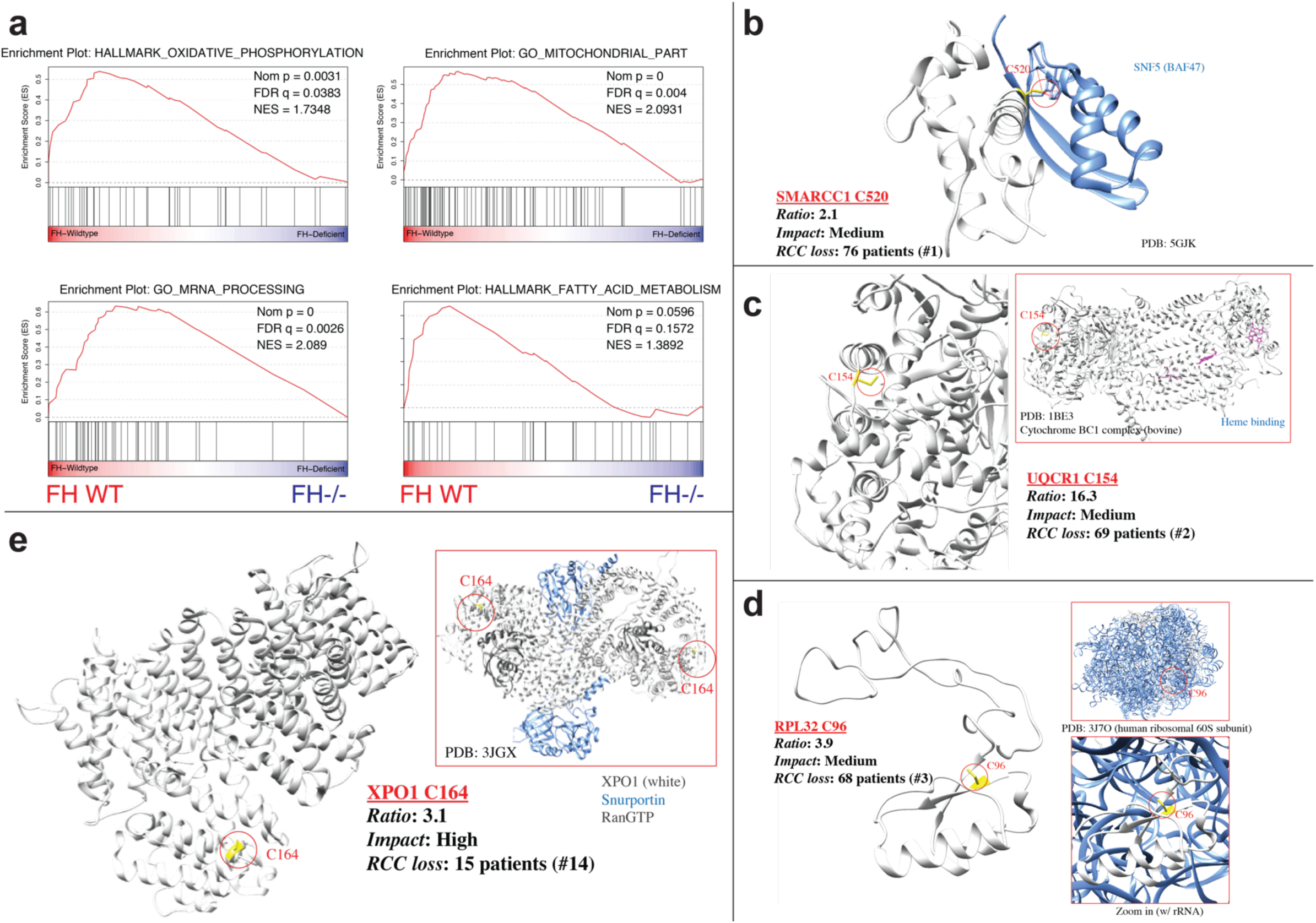
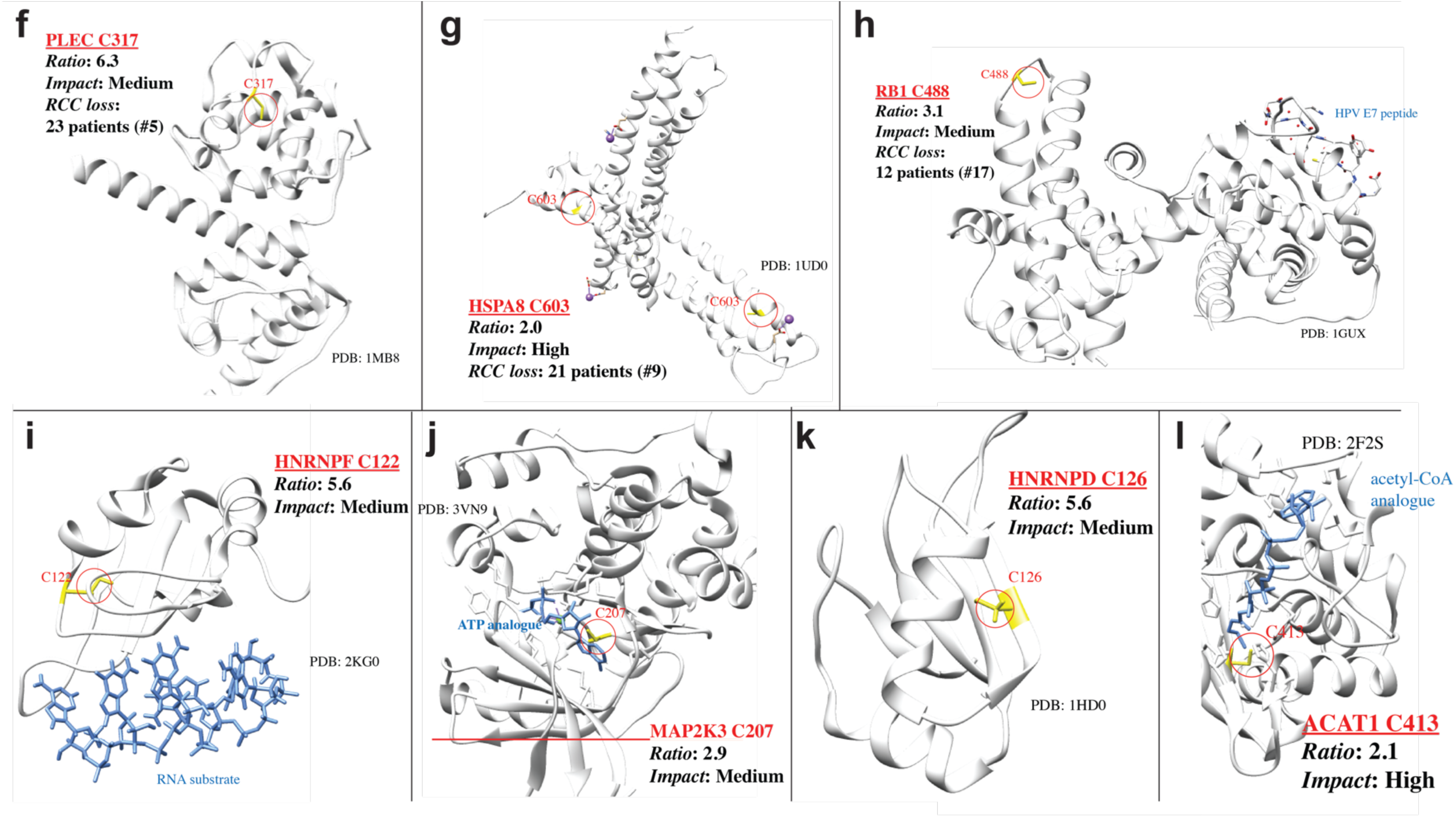
(a) Gene set enrichment analysis (GSEA) of *FH*-regulated Cys residues highlights pathways whose activity is functionally repressed in HLRCC cells. (b) Structural analysis of functional Cys residues in SMARCC1, (c) UQCR1, (d) RPL32, (e) XPO1, (f) PLEC, (g) HSPA8, (h) RB1, (i) HNRNPF, (j) MAP2K3, (k) HNRNPD, (l) ACAT1. In cases where mutations are known in kidney cancer, genes are annotated by cBioPortal “rank” (see Figure 5C, Table S4).

**Figure S5.**
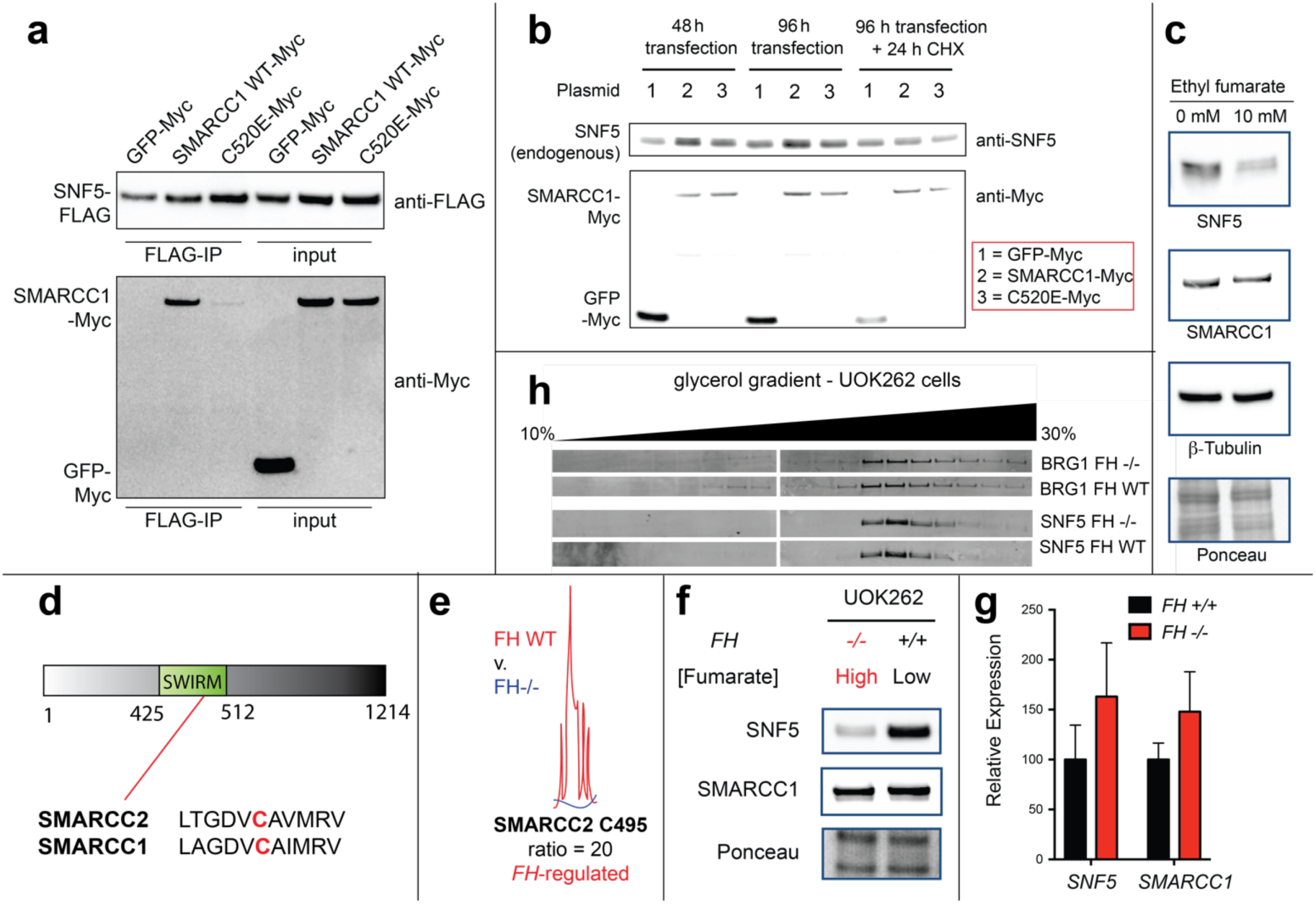
(a) Full immunoblotting data for SMARCC1-SNF5 co-immunoprecipitation in HEK-293 cells co-overexpressing FLAG-tagged SNF5 with Myc-tagged SMARCC1 (WT or C520E mutant) or control (Myc-tagged GFP). (see Figure 5g). b) Overexpression of SMARCC1-WT but not SMARCC1-C520E causes stabilization of endogenous SNF5 in HEK-293 cells compared to the GFP control. Cycloheximide (CHX, 200 μg/mL, 24 h) treatment leads to increased degradation of SNF5 in SMARCC1-overexpressing cells compared to the control. (c) Cell permeable ethyl fumarate (10 mM, 12 h treatment) causes degradation of endogenous SNF5 but not SMARCC1. (d) The SWIRM domains of SMARCC1 and SMARCC2 are highly homologous and include a conserved cysteine. (e) C495 of the SMARCC2 SWIRM (homologous to SMARCC1 C520) undergoes *FH*-regulated occupancy changes in HLRCC cells. (f) SNF5, but not SMARCC1, levels are lower in FH-deficient HLRCC cells. (g) Expression of *SNF5* and *SMARCC1* is not significantly altered by FH loss as assessed by qRT-PCR. (h) Glycerol gradient fractionation indicates *FH* status does not greatly alter SWI-SNF composition in UOK262 cell lines.

## Materials and Methods

### General materials and methods

HEK-293 cells were obtained from the NCI tumor cell repository. UOK262 (*FH*−/−), UOK262WT (*FH +/+* rescue), UOK268 (*FH*−/−) and UOK268WT (*FH +/+* rescue) cells were obtained from Linehan lab34. Plasmids encoding FLAG-tagged SNF5, Myc-tagged SMARCC1 and Myc-tagged GFP were obtained as a gift from Trevor Archer (Epigenetics & Stem Cell Biology Laboratory, NIEHS).C520E mutation was introduced to Myc-SMARCC1 entry clone using custom oligos along with the Quick Change Site-Directed Mutagenesis Kit (Agilent #200515) and transformed into DH10B cells. The insert was fully sequenced to confirm the mutation. Transfection-quality plasmid DNA was generated using the GenElute HP Maxiprep Kit. Qubit Protein Assay kit was purchased from Life Technologies (Q33212). Streptavidin agarose resin was purchased from ThermoFisher Scientific (20353). S-succinated-Cys antibody was kindly provided by Prof. Norma Frizzell (University of South Carolina). SMARCC1 (11956), SNF5 (8745), BRG1 (3508), PKM1 (7067), Myc-Tag (2278), FLAG-Tag (14793) and HA-Tag (3724) antibodies were purchased from Cell Signaling Technologies. OAT (A305-355A), HNRNP-L (A303-895A), CBX5 (A300-877A), EEF2 (A301-688A) and MAP2K4 (A302-658A) antibodies were purchased from Bethyl Laboratories, Inc. IP-grade antibodies for SMARCC1 (sc-32763) and BRG1 (sc-17796) were obtained from Santa Cruz Biotechnology. Protein A/G plus agarose resin was purchased from Sigma (20423). Fumaric acid (A10976) and ethyl fumarate (A12545) were purchased from Alfa Aesar. Maleic acid (M0375), dimethyl fumarate (242926) and mono-methyl fumarate (651419) were purchased from Sigma. Cycloheximide (14126) and EPZ6438 (16174) were purchased from Cayman chemical. Anti-FLAG pulldown was performed using immunoprecipitation kit (KBA-319-383) from Rockland Immunochemicals, Inc. SDS-PAGE was performed using Bis-Tris NuPAGE gels (4-12%, Invitrogen #NP0322), and MES running buffer (Life technologies #NP0002) in Xcell SureLock MiniCells (Invitrogen) according to the manufacturer’s instructions. SDS-PAGE fluorescence was visualized using an ImageQuant Las4010 Digitial Imaging System (GE Healthcare). Total protein content on SDS-PAGE gels was visualized by Blue-silver Coomassie stain, made according to the published procedure.^77^ For western blotting, SDS-PAGE gels were transferred to nitrocellulose membranes (Novex, Life Technologies # LC2001) by electroblotting at 30 volts for 1 hour using a XCell II Blot Module (Novex). Membranes were blocked using StartingBlock (PBS) Blocking Buffer (Thermo Scientific) for 20 minutes, then incubated overnight at 4°C in a solution containing the primary antibody of interest (1:3000 dilution for S-succinated-Cys antibody and 1:1000 dilution for all other antibodies) in the above blocking buffer with 0.05% Tween 20. The membranes were next washed with TBST buffer, and incubated with a secondary HRP-conjugated antibody (anti-rabbit IgG, HRP-linked [7074], Cell Signaling, 1:1000 dilution) for 1 hour at room temperature. The membranes were again washed with TBST, treated with chemiluminescence reagents (Western Blot Detection System, Cell Signaling) for 1 minute, and imaged for chemiluminescent signal using an ImageQuant Las4010 Digitial Imaging System (GE Healthcare).

### Cell culture and isolation of whole-cell lysates

HEK-293 cells were cultured at 37 °C under 5% CO_2_ atmosphere in a growth medium of DMEM supplemented with 10% FBS and 2 mM glutamine. UOK262 and UOK268 cell lines were cultured in DMEM supplemented with 10% FBS, 2 mM glutamine, 1 mM pyruvate. UOK262WT and UOK268WT cell lines were cultured in DMEM supplemented with 10% FBS, 2 mM glutamine, 1 mM pyruvate and 0.3 mg/mL of G418. Unfractionated proteomes were harvested from cell lines (80-90% confluency) by washing adherent cells 3x with ice cold PBS, scraping cells into a Falcon tube, and centrifuging (1400 rcf × 3 min, 4 °C) to form a cell pellet. After removal of PBS supernatant, cell pellets were either stored at −80 °C or immediately lysed by sonication. For lysis, cells were first resuspended in 1-2 mL ice cold PBS (10-20 × 10^6^ cells/mL) containing protease inhibitor cocktail (1x, EDTA-free, Cell Signaling Technology # 5871S) and PMSF (1 mM, Sigma # 78830). These samples were then lysed by sonication using a 100 W QSonica XL2000 sonicator (3 × 1s pulse, amplitude 1, 60s resting on ice between pulses). Lysates were pelleted by centrifugation (14,000 rcf × 30 min, 4 °C) and quantified on a Qubit 2.0 Fluorometer using a Qubit Protein Assay Kit. Quantified proteomes were diluted to 2 mg/mL and stored in 1 mg aliquots at −80 °C for chemoproteomic or enzyme activity analyses. For the studies involving pH-dependence, cells were lysed in a lysis buffer containing 50 mM potassium phosphate buffer at specified pH, 1 mM PMSF and 1x protease inhibitor cocktail.

### Gel-based detection of FA-alkyne labeled proteomes

20 μg proteome was incubated with specified concentration of FA-alkyne at room temperature for the specified time. For competition experiments, 1 mg proteome (0.5 mL, 2 mg/mL) was pre-incubated with the competitor (1 mM) for 3 h, followed by 15 h treatment with FA-alkyne (100 μM). Proteomes were then desalted using Illustra^TM^ NAP-5 columns(GE Healthcare # 17085301) to remove unreacted reagents and 20 μg proteomes were used for analysis. Proteins labeled by FA-alkyne were visualized by SDS-PAGE via Cu(I)-catalyzed [3 + 2] cycloaddition with a fluorescent azide as previously reported.^78-79^ Briefly, TAMRA-azide (100 μM; 5 mM stock solution in DMSO), TCEP (1 mM; 100 mM stock in H_2_O), tris-(benzyltriazolylmethyl)amine ligand (TBTA; 100 μM; 1.7 mM stock in DMSO:*tert*-butanol 1:4), and CuSO_4_ (1 mM; 50 mM stocks in H_2_O) were sequentially added to the labeled proteome. Reactions were vortexed, incubated at room temperature for 1 hour, quenched by addition of 4x SDS-loading buffer (strongly reducing) and analyzed by SDS-PAGE. Gels were fixed and destained in a solution of 50% MeOH/40% H_2_O/10% AcOH overnight to remove excess probe fluorescence, rehydrated with water, and visualized using an ImageQuant Las4010 (GE Healthcare) with green LED excitation (λ_max_ 520–550 nm) and a 575DF20 filter.

### Chemoproteomic labeling and enrichment of fumarate-sensitive and *FH*-regulated cysteines

For profiling of fumarate-sensitive cysteines, 2 mg of HEK-293 proteomes (1 mL, 2 mg/mL) were incubated with 1 mM fumaric acid (10 μL, 100 mM stock in DMSO) or vehicle (DMSO, 10 μL) overnight at room temperature, followed by labeling with 100 μM IA-alkyne (10 μL, 10 mM stock in DMSO) for 1 h. Proteomes were then desalted using NAP-5 columns to remove unreacted reagents. For identification of *FH*-regulated cysteines, 2 mg of UOK262 or UOK262WT proteomes (1 mL, 2 mg/mL) were labeled with 100 µM IA-alkyne (10 μL, 10 mM stock in DMSO) for 1 h at room temperature. For enrichment of fumarate-sensitive and *FH*-regulated cysteines, probe labeled proteins were then conjugated to light (low fumarate proteomes: vehicle-treated HEK-293 or UOK262WT) or heavy (high fumarate proteomes: fumarate-treated HEK-293 or UOK262) diazobenzene biotin-azide (azo) tag by Cu(I)-catalyzed [3 + 2] cycloaddition as previously reported.^80^ Briefly, azo-tag (100 μM), TCEP (1 mM), TBTA (100 μM), and CuSO_4_ (1 mM) were sequentially added to the labeled proteome. Reactions were vortexed and incubated at room temperature for 1h. Proteomes labeled with heavy and light azo-tags were then combined pairwise and centrifuged (6500 rcf *x* 10 min, 4 °C) to collect precipitated protein. Supernatant was discarded, and protein pellets were resuspended in 500 μL of methanol (dry-ice chilled) with sonication, and centrifuged (6500 rcf *x* 10 min, 4 °C). This step was repeated, and the resulting washed pellet was redissolved (1.2% w/v SDS in PBS; 1 mL); sonication followed by heating at 80-95 °C for 5 min was used to ensure complete solubilization. Samples were cooled to room temperature, diluted with PBS (5.9 mL), and incubated with Streptavidin beads (100 μL of 50% aqueous slurry per enrichment) overnight at 4 °C. Samples were allowed to warm to room temperature, pelleted by centrifugation (1400 rcf *x* 3 min), and supernatant discarded. Beads were then sequentially washed with 0.2% SDS in PBS (5 mL × 1), PBS (5 mL × 3) and H_2_O (5 mL × 3) for a total of 7 washes.

### On bead reductive alkylation, tryptic digest and diazobenzene cleavage of proteomic samples

Following the final wash, protein-bound streptavidin beads were resuspended 6 M urea in PBS (500 μL) and reductively alkylated by sequential addition of 10 mM DTT (25 μL of 200 mM in H_2_O, 65 °C for 20 min) and 20 mM iodoacetamide (25 μL of 400 mM in H_2_O, 37 °C for 30 min) to each sample. Reactions were then diluted by addition of PBS (950 μL), pelleted by centrifugation (1400 rcf × 3 min), and the supernatant discarded. Samples were then subjected to tryptic digest by addition of 200 μL of a pre-mixed solution of 2M urea in PBS, 1 mM CaCl_2_ (2 μL of 100 mM in H_2_O), and 2 μg of Trypsin Gold (Promega, 4 μL of 0.5 μg/μL in 1% acetic acid). Samples were shaken overnight at 37 °C and pelleted by centrifugation (1400 rcf × 3 min). Beads were then washed sequentially with PBS (500 µL × 3) and H_2_O (500 µL × 3). Labeled peptides were eluted from the beads by sodium dithionite mediated cleavage of the diazobenze of the azo-tag. For this, beads were incubated with freshly prepared 50 mM sodium dithionite in PBS (50 µL) for 1 h at room temperature. Beads were pelleted by centrifugation (1400 rcf × 3 min) and supernatant was transferred to a new Eppendorf tube. The cleavage process was repeated twice more with 50 mM sodium dithionite (75 µL) and supernatants were combined with the previous. The beads were additionally washed two times with water (75 µL) and supernatants were collected and combined with previous. Formic acid (17.5 µL) was added to the combined supernatants and samples were stored at −20 °C until ready for LC-MS/MS analysis.

### LC-MS/MS and data analysis for quantitative cysteine reactivity profiling

Mass spectrometry was performed using a Thermo LTQ Orbitrap Discovery mass spectrometer coupled to an Agilent 1200 series HPLC. Labeled peptide samples were pressure loaded onto 250 mm fused silica desalting column packed with 4 cm of Aqua C18 reverse phase resin (Phenomenex). Peptides were eluted onto a 100 mm fused silica biphasic column packed with 10 cm C18 resin and 4 cm Partisphere strong cation exchange resin (SCX, Whatman), using a five step multidimensional LC-MS protocol (MudPIT). Each of the five steps used a salt push (0%, 50%, 80%, 100%, and 100%), followed by a gradient of buffer B in Buffer A (Buffer A: 95% water, 5% acetonitrile, 0.1% formic acid; Buffer B: 20% water, 80% acetonitrile, 0.1% formic acid) as outlined previously.^81-82^ The flow rate through the column was ∼0.25 µL/min, with a spray voltage of 2.75 kV. One full MS1 scan (400-1800 MW) was followed by 8 data dependent scans of the n^th^ most intense ion. Dynamic exclusion was enabled. The tandem MS data, generated from the 5 MudPIT runs, was analyzed by the SEQUEST algorithm.^83^ Static modification of cysteine residues (+57.0215 m/z, iodoacetamide alkylation) was assumed with no enzyme specificity. The precursor-ion mass tolerance was set at 50 ppm while the fragment-ion mass tolerance was set to 0 (default setting). Data was searched against a human reverse-concatenated non-redundant FASTA database containing Uniprot identifiers. MS datasets were independently searched with light and heavy azo-tag parameter files; for these searches differential modifications on cysteine of +456.2849 (light) or +462.2987 (heavy) were used. MS2 spectra matches were assembled into protein identifications and filtered using DTASelect2.0,^84^ with the –trypstat and –modstat options applied. Peptides were restricted to fully tryptic (-y 2) with a found modification (-m 0) and a delta-CN score greater than 0.06 (-d 0.06). Single peptides per locus were also allowed (-p 1) as were redundant peptides identifications from multiple proteins, but the database contained only a single consensus splice variant for each protein. RAW files have been uploaded to the PRIDE database are directly available for the authors during review upon request. Quantification of L/H ratios were calculated using the cimage quantification package described previously.^26^

### Whole proteome protein abundance analysis

100 µg of UOK262 or UOK262WT proteomes (100 µL, 1 mg/mL) were precipitated by the addition of 5 µL 100% trichloroacetic acid in PBS, vortexed and frozen at −80 °C overnight. Samples were thawed, and proteins were pelleted by centrifugation (17000 rcf × 10 min). Each protein pellet was washed by resuspension in acetone (500 µL) using sonication, followed by centrifugation (2200 rcf × 10 min). Supernatant was discarded, pellet was allowed to dry and then resuspended thoroughly by sonication in 30 µL 8M urea in PBS. Reductive alkylation was then performed by sequential addition of 70 µL of 100 mM ammonium bicarbonate and 1.5 µL of 1M DTT (65 °C for 15 min) and iodoacetamide (2.5 μL of 400 μM in H_2_O, room temperature for 30 min). Reactions were then diluted by addition of PBS (120 μL) and tryptic digest was performed by addition of 2 μg of Trypsin Gold and and 2.5 µL of 100 mM CaCl_2_, followed by overnight incubation at 37 °C. Trypsin was quenched by addition of 10 µL formic acid (∼5% final volume) and undigested protein was pelleted by centrifugation (17000 rcf × 20 min). Supernatant was collected and stored at −20 °C until ready for LC-MS/MS analysis which was performed using ∼50 µL of each sample. LC-MS/MS was performed as described above with slight modification to MudPIT protocol. Here salt pushes of 0%, 25%, 50%, 80%, and 100% were employed. Tandem MS data analysis was performed as described above. Spectral counting was used for calculating the UOK262WT:UOK262 protein abundance ratios for those proteins which had >10 spectral counts in at least one of the two cell lines and these ratios were used to correct the *FH*-regulated cysteine ratios wherever possible.

### Bioinformatic analysis of fumarate-sensitive and *FH*-regulated cysteines

Annotation of protein subcellular localization as well as cysteine function and conservation was generated from the Uniprot Protein Knowledgebase (UniProtKB) as described previously.^33, 85^ Analysis of linear sequences flanking fumarate-sensitive and *FH*-regulated cysteines was performed using the informatics tool pLogo, accessible at: https://plogo.uconn.edu. Input sequences are listed in Table S4, and were derived from the 50 cysteines found to be most fumarate-sensitive and insensitive in this study, (highest and lowest R values, n≥2, SD≤25%), as well as the 50 cysteines found to be most hyperreactive,^26^ DMF-sensitive,^30^ MMF-sensitive,^30^ GSNO,^32^ and HNE-sensitive^37^ in literature datasets. Protein sequences for motif analysis were derived from their tryptic peptide sequences using Peptide Extender (schwartzlab.uconn.edu/pepextend). Conservation and functional impact of fumarate-sensitive and *FH*-regulated cysteines identified in chemoproteomic experiments was analyzed using the informatics tool Mutation Assessor, accessible at: http://mutationassessor.org/r3. Conservation analysis depicted in Fig. 2d represents the output of the variant conservation (VC) score for the 50 cysteines found to be most fumarate-sensitive in this study (highest L/H ratio [R] values, n≥2, SD≤25%).), the 50 cysteines found to be least fumarate-sensitive in this study (lowest R values), and the 50 cysteines found to be most hyperreactive in a previous chemoproteomic study performed by Weerapana et al.^26^ Potential functional impact of fumarate modifications (Fig. S5a, Table S4) reflects the effect of C to E mutations on the functional impact (FI) output of Mutation Assessor. Gene ontology analysis was performed using the bioinformatics tool DAVID, accessible at: http://david.ncifcrf.gov/. Output tables in Table S4 reflect DAVID analysis of fumarate-sensitive and *FH*-regulated cysteines predicted to have a medium or high impact on protein function by Mutation Assessor. Candidate functional fumarate targets were assessed for cases of genomic alteration in renal cell carcinoma (clear cell and non-clear cell) using cBioPortal (http://cbioportal.org). Structural analysis of candidate functional fumarate targets known to undergo genomic alteration in renal cell carcinoma was performed using Chimera. For gene set enrichment analysis (GSEA, Fig. 6), R values for fumarate-sensitive and *FH*-regulated peptides were log2-transformed and analyzed for 1000 permutations using the Broad Institute’s javaGSEA desktop application (http://software.broadinstitute.org/gsea/downloads.jsp). For proteins in which R values were measured for more than one cysteine-containing peptide, the peptide with the greatest absolute R value was used for GSEA analysis. GSEA outputs were re-plotted for graphics using a variant of ReplotGSEA package, accessible at: https://github.com/PeeperLab/Rtoolbox/blob/master/R/ReplotGSEA.R.

### Validation of fumarate-sensitive and *FH*-regulated targets using FA-alkyne

5 mg of HEK-293 proteome (2.5 mL, 2 mg/mL) was pre-treated with 1 mM fumaric acid (25 μL, 100 mM stock in DMSO) or DMSO for 3 hours prior to incubation with 100 μM FA-alkyne (25 μL, 10 mM stock in DMSO) for 15 hours. Proteomes were then desalted using Illustra^TM^ NAP-25 columns(GE Healthcare # 17085201) to remove unreacted reagents. Labeled proteomes were enriched via Cu(I)-catalyzed [3 + 2] cycloaddition with biotin-azide as described above for chemoproteomic analysis. Following the final wash, enriched resin was collected on top of centrifugal filters (VWR, 82031-256). Proteins were eluted from resin via addition of 40 μL 1x SDS sample buffer, followed by boiling for 10 min at 95 °C. Following repetition of the elution step, both eluents were combined and 20 μL of the combined eluent was loaded onto a 4-12% SDS-PAGE gel and analyzed by western blotting.

### Fluorescent quantification of fumarate release from *S*-succinated thiols

*S*-succinated thiols (1 mM final concentration, 5 μL of 20 mM stock in DMSO) were incubated in TRIS buffer (100 mM; pH 7, 7.5, 8, and 8.5) at 37 °C for 24 h. After incubation, reactions were developed by treatment with equal volume of hydrazonyl chloride **4** from Zengeya et al.^38^ (150 μM final concentration, 300 μM stock in CH_3_CN) for 1 h at room temperature. Fluorescence produced was then measured on Photon Technology International QuantMaster fluorimeter using 1-cm path length, 0.13 mL quartz microcuvettes (Helma #101-015-40) at ambient temperature (22 ± 2 °C), using an excitation wavelength of 390 nm, slit width of 3.5 nm, and monitoring emission from 410 nm to 615 nm. Fluorescence emission values at 530 nm were used to calculate percent DMF released by interpolating into a standard curve of DMF reacting hydrazonyl chloride **4** under identical conditions.

### Ectopic expression and co-immunoprecipitation of SMARCC1 and SNF5

HEK-293 cells were plated in 10 cm dishes (3×10^6^ cells/dish in 10 ml DMEM media/well), and allowed to adhere and grow for 24 h. FLAG-tagged SNF5 was co-overexpressed with Myc-tagged GFP, SMARCC1 or SMARCC1-C520E using lipofectamine 2000 (Invitrogen # 11668019) according to the manufacturer’s instructions. Co-overexpressions were carried out by incubating the cells for 48 h at 37 °C under 5% CO_2_ atmosphere, after which the cells were harvested, soluble proteome isolated and quantified as described above. Anti-FLAG pulldown was performed using immunoprecipitation kit (KBA-319-383) according to the manufacturer’s instruction. 1 mg of the lysate was incubated with the anti-FLAG resin overnight at 4 °C. Purified protein was ran on SDS-PAGE and immunoblotted against anti-Myc-tag and anti-FLAG-tag.

### Cellular analysis of SMARCC1 overexpression on SNF5 levels

HEK-293 cells were plated in 6-well dishes (6×10^5^ cells/well in 3 ml DMEM media/well), and allowed to adhere and grow for 24 h. At this point, transient transfection of plasmids encoding for Myc-tagged GFP, SMARCC1 or SMARCC1(C520E) was performed using lipofectamine 2000 (Invitrogen # 11668019) according to the manufacturer’s instructions. Overexpression was carried out by incubating the cells for 48 h at 37 °C under 5% CO_2_ atmosphere. For the cycloheximide treatment experiment, overexpression was carried out for 96 h. After 96 h, media was changed and cells were incubated with 200 μg/mL cycloheximide or vehicle for additional 24 h. After the treatment, cells were harvested, and soluble proteome was isolated and quantified as described above. 10 μg of lysates were loaded per lane of the gel for the western blot analysis of endogenous SNF5 and expression levels of Myc-tagged GFP, SMARCC1 or SMARCC1(C520E).

### Co-immunoprecipitation of endogenous SMARCC1 and BRG1 in HLRCC cells

For co-immunoprecipitation of endogenous SMARCC1 and BRG1, whole cell lysates from HLRCC cells were first prepared by resuspending cell pellets in IP-buffer containing 50 mM Tris pH 8, 400 mM NaCl, 2 mM EDTA, 10% glycerol, 1% NP-40 (Ipegal® CA-630, Sigma # I8896), 1 mM PMSF and 1X protease inhibitor cocktail. The lysates were pelleted by centrifugation (14,000 rcf × 30 min, 4 °C) and pre-cleared by incubating with protein A/G plus agarose resin (30 μL) for 1 h at 4 °C. Pre-cleared supernatant was collected by centrifugation (10,000 rcf × 5 min, 4 °C) and diluted to 1 mg/mL concentration. For each co-immunoprecipitation, 2 mg of whole cell proteome was incubated with 2.5 μg/mL of SMARCC1 (sc-32763) or BRG1 (sc-17796) antibody at 4 °C for 1 h. Protein A/G plus agarose resin (100 μL) was added to each sample and incubated overnight at 4 °C. Samples were pelleted by centrifugation, supernatant was discarded, and beads were then washed with IP-buffer (1 mL × 3). Enriched proteins were eluted from resin via addition of 40 μL 1× SDS sample buffer, followed by boiling for 10 min at 95 °C. Following repetition of the elution step, both eluents were combined and 20 μL of the combined eluent was loaded onto a 4-12% SDS-PAGE gel and analyzed by western blotting.

### qRT-PCR analysis of SWI/SNF expression in HLRCC cells

Total RNA was isolated from 1 × 10^6^ cells using the RNeasy Plus mini kit (Qiagen cat. #74136) and 500-1000 ng of purified RNA was used as template for cDNA synthesis (Life Technologies, cat. #18080-051). mRNA expression of *SMARCC1* and *SNF5* in UOK262 cells was normalized to the housekeeping gene *ACTB* (β-actin). *SMARCC1* and *SNF5* primers were taken from DelBove et al.^47^, while *ACTB* primers were taken from PrimerBank.^86^ Primers used were as follows-*SMARCC1*: CACCCCAGCCAGGTCAGAT (forward) and TGCAACAGTGGGAATCATGC (reverse); *SNF5*: CAGAAGACCTACGCCTTCAG (forward) and GTCCGCATCGCCCGTGTT (reverse); and *ACTB*: CATGTACGTTGCTATCCAGGC (forward) and CTCCTTAATGTCACGCACGAT (reverse). Quantitative real-time PCR was performed using PerfeCTa SYBR Green FastMix (Quanta Biosciences, cat. #95072) and a mastercycler ep-gradient realplex^2^ (Eppendorf). Baseline was set manually at two cycles prior to the earliest visible amplification of fluorescence. Threshold was set manually at the lower half of linear phase amplification. Results are expressed as fold change above UOK262WT cells after normalization to *ACTB* expression using the ΔΔCt method, as previously described.^87^

### Inhibition of HLRCC spheroid growth by EZH2 inhibitors

For tumor spheroid formation, a total of 5000 single cell suspensions were plated in 100 μL of complete media (DMEM supplemented with 20% FBS, 1x MEM non-essential amino acids, and 1x Anti-Anti) into each well of a 96-well ultra-low attachment plates (Corning 3603). After 3 days in culture, tumor spheroid formation was confirmed visually using the EVOS XL Core Cell Imaging System (Thermo Fisher Scientific). On day 0, 100 μL of media containing 2x concentration of EPZ6438 was added to the wells diluting the compound to the indicated concentration. Every 3 or 4 days, 100 μL of media was removed and replaced with 100 μL of media with 2x concentration of EPZ6438. The spheroids were treated for 21 days. The spheroids where then dissociated with Cell Titer Glo 3D (Promega # G9681) following manufacturer’s instructions. The plates were then read on an Enspire Mulitmode Plate Reader (Perkin-Elmer).

### Analysis of SWI/SNF complex composition by glycerol gradient fractionation in HLRCC cells

UOK262 *FH* −/− and *FH+/+* rescue cells were grown to 90% confluency in 2 × 15 cm dishes per cell line. Cells were harvested by trypsinization and washed once in ice-cold PBS. Nuclei were isolated by incubating the cell pellets in Buffer A (20 mM HEPES pH 7.9, 25 mM KCl, 10% glycerol, 0.1% NP-40, 1 mM DTT with PMSF, aprotinin, leupeptin and pepstatin) for 7 min. Nuclei were pelleted and washed in buffer A without NP-40. Washed and pelleted nuclei were resuspended in Buffer C (10 mM HEPES, pH 7.6, 3 mM MgCl_2_, 100 mM KCl, 0.1 mM EDTA, 10% glycerol, 1 mM DTT with PMSF, aprotinin, leupeptin and pepstatin). Ammonium sulfate was added to 0.3 M final concentration. Samples were incubated in a rotating wheel at 4 °C for 30 min and cleared by ultracentrifugation (150000 rcf × 30 min). 300 mg of ammonium sulfate powder was introduced per mL of cleared lysate. After ice incubation for 20 min, proteins were precipitated by ultracentrifugation (150000 rcf × 30 min). Pelleted proteins were resuspended in 100 μL HEMG1000 buffer (25 mM HEPES pH 7.6, 0.1 mM EDTA, 12.5 mM MgCl_2_, 100 mM KCl, 1 mM DTT with PMSF, aprotinin, leupeptin and pepstatin). 400 μg of resuspended proteins were layered over 10 mL, 10-30% glycerol gradient, prepared with HMG1000 buffer without glycerol or with 30% glycerol, and separated by centrifugation at 40000 rpm (Beckman Coulter XL-100K, Brea, CA) for 16 h using SW32Ti rotor (Beckman Coulter, Brea, CA). 500 μL-fractions were collected and analyzed by western blotting using antibodies against BRG1 (Abcam, ab110641) and SNF5 (Santa Cruz Biotechnology, sc-166165).

## Contributions

R.A.K., D.W.B., and J.L.M. designed experiments. R.A.K. and D.W.B. performed all chemical proteomic labeling and enrichment experiments. D.W.B. performed all LC-MS/MS studies and stoichiometry analyses of IA-alkyne enrichment experiments. R.A.K. and A.B. synthesized compounds. R.A.K. and S.B. performed all cell-based analyses. C.B. assisted with cell-based analyses and performed S-succination reversibility studies. J.H.S. performed co-immunoprecipitation experiments. J.L.M., D.B., A.T., and R.A.K. performed bioinformatics analyses. D.W. and W.M.L. performed HLRCC spheroid growth inhibition studies and assisted with SWI/SNF analyses. A.A. and E.C.D. performed glycerol gradient fractionation and analysis of SWI/SNF complex in HLRCC cell lines. N.F. provided the S-succination antibody. W.M.L. provided HLRCC cell lines and advised experimental design. M.P.W., L.F., and M.L. performed whole proteome MUDPIT analyses of HLRCC cells for S-succination validation. R.A.K. and J.L.M. analyzed data and wrote the manuscript with input from all authors.

## Acknowledgements

The authors thank Dr. Carissa Grose (Protein Expression Laboratory) for assisting with cloning and preparation of plasmid DNA, Dr. Trevor Archer (Epigenetics & Stem Cell Biology Laboratory, NIEHS) for the kind gift of the SMARCC1 and SNF5 plasmids, Allison Roberts, Julie Garlick, and Dr. Thomas Zengeya (Chemical Biology Laboratory, NCI) for assisting with preliminary studies, and Dr. Dan Crooks and Dr. Chris Ricketts (Urologic Oncology Branch, NCI) for helpful discussions. This work was supported by the Intramural Research Program of the NIH, National Cancer Institute, Center for Cancer Research (ZIA BC011488-04).

