## Supplementary Materials for "A chemoproteomic portrait of the oncometabolite fumarate"

**Supplemental Information**

**MudPIT** **analysis of HLRCC proteomes for validation of S-succination of fumarate-sensitive and *FH*-regulated cysteine residues**

TCA-precipitated protein samples from whole cell extracts from UOK262 and UOK268 *FH -/-* cells were analyzed independently in triplicate by Multidimensional Protein Identification Technology (MudPIT), as described previously.^1-2^ After recombinant endoproteinase LysC and trypsin digestions, peptide mixtures were pressure-loaded onto 100 µm fused silica microcapillary columns packed first with 9 cm of reverse phase material (Aqua; Phenomenex), followed by 3 cm of 5-μm Strong Cation Exchange material (Luna; Phenomenex), followed by 1 cm of 5-μm C_18_ RP. The loaded microcapillary columns were placed in-line with a 1260 Quartenary HPLC (Agilent). The application of a 2.5 kV distal voltage electrosprayed the eluting peptides directly into Orbitrap-Velos Pro or Elite hybrid mass spectrometers (Thermo Scientific) equipped with a custom-made nano-LC electrospray ionization source. Full MS spectra were recorded on the eluting peptides over a 400 to 1600 *m*/*z* range in the Orbitrap at 60K resolution, followed by fragmentation in the ion trap (at 35% collision energy) on the first to fifteenth most intense ions selected from the full MS spectrum. Dynamic exclusion was enabled for 90 sec.^3^ Mass spectrometer scan functions and HPLC solvent gradients were controlled by the XCalibur data system (Thermo Scientific).

RAW files were extracted into .ms2 file format^4^ using RawDistiller v. 1.0, in-house developed software.^5^ RawDistiller D(g, 6) settings were used to abstract MS1 scan profiles by Gaussian fitting and to implement dynamic offline lock mass using six background polydimethylcyclosiloxane ions as internal calibrants.^5^ MS/MS spectra were first searched using ProLuCID^6^ with a peptide mass tolerance of 10 ppm and 500 ppm for fragment ions. Trypsin specificity was imposed on both ends of candidate peptides during the search against a protein database combining 36,628 human proteins (NCBI 2016-06-10 release), as well as 193 usual contaminants such as human keratins, IgGs and proteolytic enzymes. To estimate false discovery rates (FDR), each protein sequence was randomized (keeping the same amino acid composition and length) and the resulting "shuffled" sequences were added to the database, for a total search space of 73,642 amino acid sequences. Masses of 57.0215 Da and 116.0112 Da were differentially added to cysteine residues to account for alkylation by CAM and succination, respectively, while 15.9949 Da were differentially added to methionine residues.

DTASelect v.1.9^7^ was used to select and sort peptide/spectrum matches (PSMs) passing the following criteria set: PSMs were only retained if they had a DeltCn of at least 0.08; minimum XCorr values of 1.8 for singly-, 2.0 for doubly-, and 3.0 for triply-charged spectra; peptides had to be at least 7 amino acids long. Results from each sample were merged and compared using CONTRAST.^7^ Combining all six runs, proteins had to be detected by at least 2 peptides and/or 4 spectral counts. Proteins that were subsets of others were removed using the parsimony option in DTASelect on the proteins detected after merging all runs. Proteins that were identified by the same set of peptides (including at least one peptide unique to such protein group to distinguish between isoforms) were grouped together, and one accession number was arbitrarily considered as representative of each protein group.

*NSAF7*^8^ was used to create the final reports on all detected peptides and non-redundant proteins identified across the different runs. Spectral and protein level FDRs were, on average, 0.29 ± 0.04% and 2.7 ± 0.4%, respectively. *NSAF7* was also used to generate a list of all peptide to spectrum matches (PSMs) leading to the identification of succinylated proteins. *NSAF7* was used to create PDF files displaying fully annotated MS/MS spectra (shown below) matching the modified peptides listed in Supporting Table S5.

***Data accessibility***

The complete mass spectrometry datasets (raw, peak, ProLuCID search, as well as DTASelect result, and protein sequences fasta files) for UOK262 MudPIT analyses may be obtained via ftp://massive.ucsd.edu/MSVXXX with password (RAK20712).

***Spectra for novel S-succinated peptides***

*GCLM*

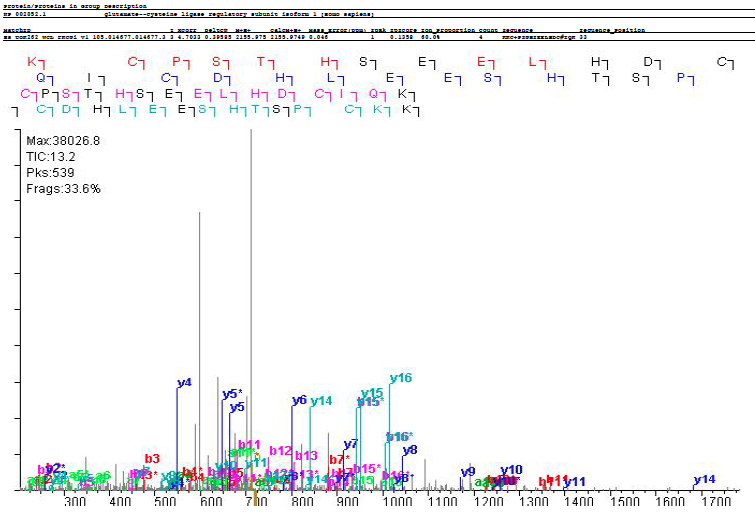

*GCLM*

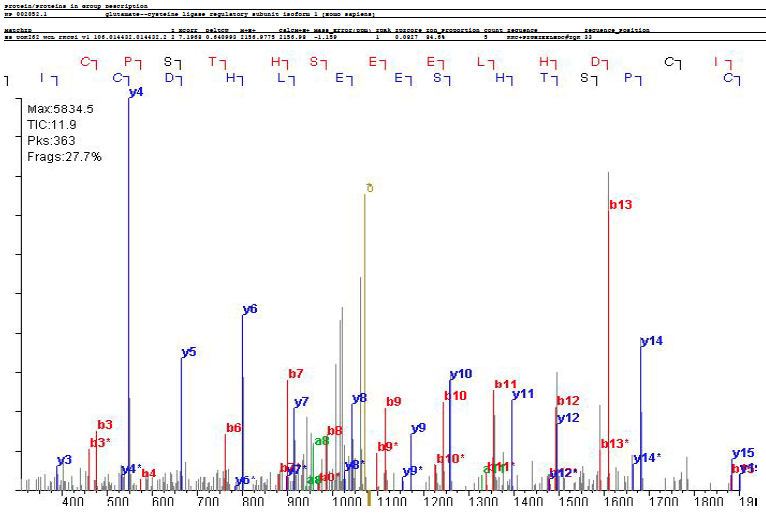

*PCBP1*

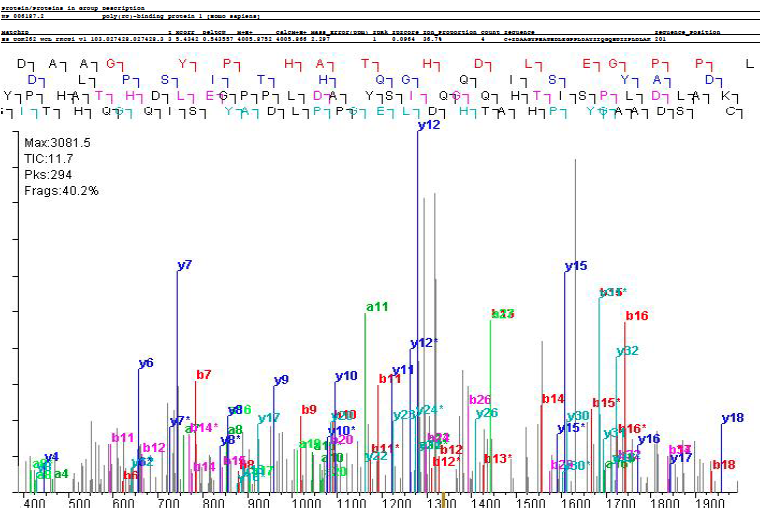

*PCBP1*

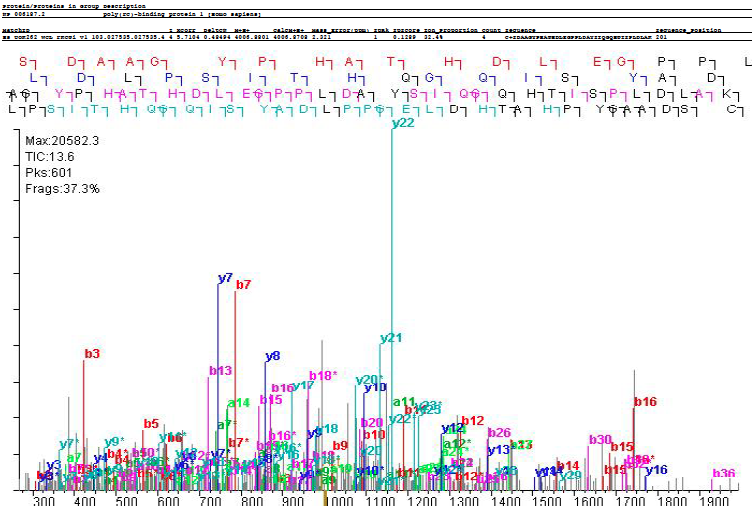

*TCP-1*

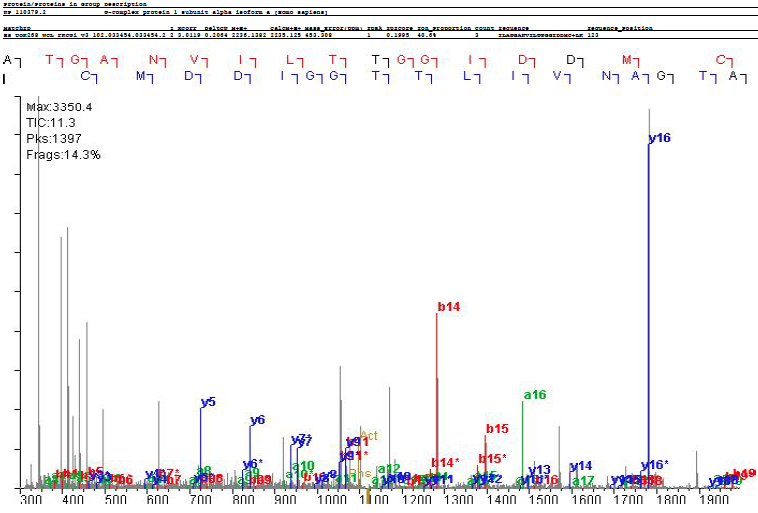

**General synthetic procedures**

Chemicals were purchased from commercial sources (Sigma-Aldrich, Alfa Aesar, and TCI America) and used without further purification unless otherwise noted. Thin-layer chromatography (TLC) was conducted with E. Merck silica gel 60 F254 precoated plates (0.25 mm) and visualized by exposure to UV light (254 nm) or chemical staining. Flash chromatography was performed using normal or reverse phase on a CombiFlash® Rf 200i (Teledyne Isco Inc). ^1^H NMR spectra were recorded at 400 MHz, and are reported relative to deuterated solvent signals. Data for ^1^H NMR spectra are reported as follows: chemical shift (δ ppm), multiplicity, coupling constant (Hz), and integration. ^13^C NMR spectra were recorded at 100 MHz, and are reported in terms of chemical shift (δ ppm). All NMR spectra were standardized to the NMR solvent signal, as specified by Gottlieb and coworkers.^9^ Analytical LC/MS was performed using a Shimadzu LC/MS-2020 Single Quadrupole utilizing a Kinetex 2.6 μm C18 100 Å (2.1 x 50 mm) column obtained from Phenomenex Inc. Runs employed a gradient of 0→90% MeCN/0.1% aqueous formic acid over 4 minutes at a flow rate of 0.2 mL/min. High-resolution LC/MS analyses were conducted on a Thermo-Fisher LTQ-Orbitrap-XL hybrid mass spectrometer system with an Ion MAX API electrospray ion source in positive or negative ion mode.

**Synthesis of fumarate alkyne (FA-alkyne)**

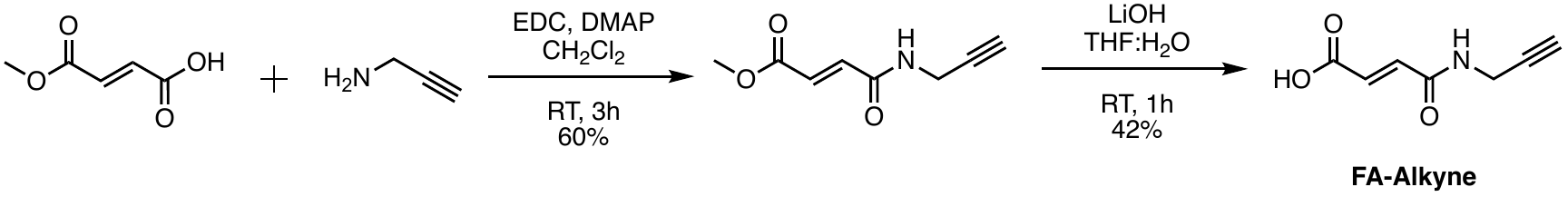

(a) **Methyl (*E*)-4-oxo-4-(propylnylamino)-2-butenoate**

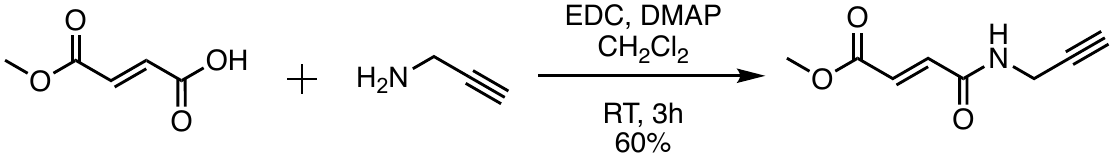

1-Ethyl-3-(3-dimethylaminopropyl)carbodiimide (0.383 g, 2.0 mmol), propargylamine (0.141 mL, 2.2 mmol) and catalytic amount of 4-dimethylaminopyridine were added to a stirring solution of mono-methyl fumarate (0.26 g, 2.0 mmol) in CH_2_Cl_2_ (10 mL). The reaction mixture was stirred at room temperature and monitored by TLC. After 3 h, the reaction mixture was diluted with CH_2_Cl_2_ (20 mL), and washed with brine (3 x 20 mL). The organic layer was dried over anhydrous Na_2_SO_4_, filtered, and concentrated under reduced pressure. This crude mixture was purified by flash chromatography (0🡪50% CH_3_OH:CH_2_Cl_2_) to yield the product **2** as yellow solid (60%). ^1^H NMR (400 MHz, CDCl_3_) δ 6.95 (d, *J* = 16.0 Hz, 1H), 6.86 (d, *J* = 16.0 Hz, 1H), 6.26 (s, 1H), 4.16 (dd, *J* = 5.3, 2.6 Hz, 2H), 3.80 (s, 3H), 2.27 (t, *J* = 2.6 Hz, 1H). ^13^C NMR (100 MHz, CDCl_3_) δ 166.08, 163.29, 135.76, 130.90, 78.80, 72.39, 52.45, 29.77. MS (ESI+): 168.2 (M+H^+^).

(b) **(*E*)-4-oxo-4-(propylnylamino)-2-butenoate**

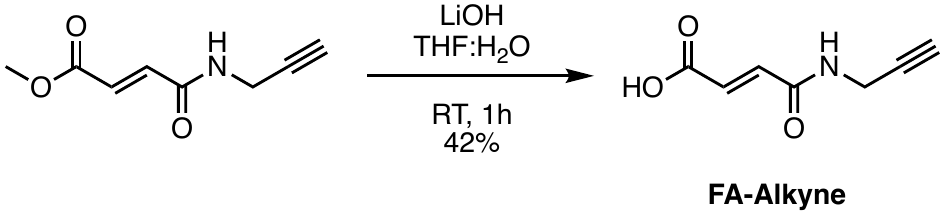

Lithium hydroxide (0.025 g, 0.598 mmol) was added to a stirring solution of **2** (0.100 g, 0.598 mmol) in a 1:1 mixture of water and tetrahydrofuran (5 mL). The reaction mixture was stirred at room temperature and monitored by TLC. After 1 h, the reaction mixture was acidified with 1M HCl (1 mL), and the crude mixture was purified by HPLC. HPLC purification was performed on a Waters HPLC with a 2545 pump and a Phenomenex Luna 10 micron C18 column (75 x 30 mm), using 0🡪95% acetonitrile/water gradient with 0.1% TFA over 10 minutes at a flow rate of 36 mL/min. Fractions containing pure compound were lyophilized to yield the product **1** as white solid (42%). ^1^H NMR (400 MHz, DMSO-*d*_6_) δ 8.98 (t, *J* = 5.5 Hz, 1H), 6.91 (d, *J* = 15.5 Hz, 1H), 6.55 (d, *J* = 15.5 Hz, 1H), 3.96 (dd, *J* = 5.4, 2.5 Hz, 2H), 3.17 (t, *J* = 2.5 Hz, 1H). ^13^C NMR (100 MHz, DMSO-*d*_6_) δ 166.34, 162.90, 136.02, 130.43, 80.41, 73.48, 28.15. MS (ESI+): 154.2 (M+H^+^).

**4-Oxo-4-(propylnylamino)butanoic acid**

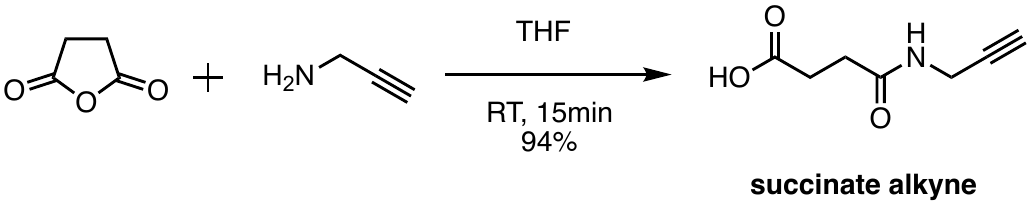
**­**

Succinic anhydride (0.200 g, 2.0 mmol) was added to a stirring solution of propargylamine (0.141 mL, 2.2 mmol) in tetrahydrofuran (5 mL). The reaction mixture was stirred at room temperature and monitored by TLC. The reaction went to completion in 15 min. At this point the crude mixture was purified by flash chromatography (0🡪20% CH_3_OH:CH_2_Cl_2_) to yield the product **1** as white solid (94%). ^1^H NMR (400 MHz, DMSO-*d*_6_) δ 12.06 (s, 1H), 8.28 (t, *J* = 5.6 Hz, 1H), 3.84 (dd, *J* = 5.5, 2.6 Hz, 2H), 3.08 (t, *J* = 2.5 Hz, 1H), 2.42 (dd, *J* = 7.1 5.8 Hz, 2H), 2.32 (dd, *J* = 7.1 5.8 Hz, 2H). ^13^C NMR (100 MHz, CD_3_OD) δ 176.16, 174.10, 80.56, 72.10, 31.29, 20.09, 29.46. MS (ESI+): 156.1 (M+H^+^).

**Dimethyl 2-(phenylthio)succinate**

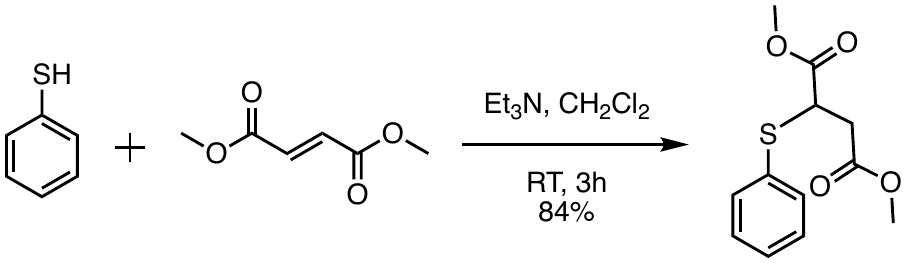

Dimethyl fumarate (0.173 g, 1.2 mmol) and triethylamine (0.167 mL, 1.2 mL) were added to a stirring solution of thiophenol (0.102 mL, 1.0 mmol) in dichloromethane (5 mL). The reaction mixture was stirred at room temperature and monitored by TLC. After 3 h, the reaction mixture was concentrated under reduced pressure and purified by flash chromatography (0🡪50% EtOAc:hexanes) to yield the product as colorless liquid (84%). ^1^H NMR (400 MHz, CDCl_3_) δ 7.55-7.43 (m, 2H), 7.39-7.29 (m, 2H), 4.02 (dd, *J* = 9.6, 5.7 Hz, 1H), 3.71 (s, 3H), 3.68 (s, 3H), 2.96 (dd, *J* = 17.0, 9.6 Hz, 1H), 2.75 (dd, *J* = 17.0, 5.7 Hz, 1H). ^13^C NMR (100 MHz, CDCl_3_) δ 171.68, 171.18, 134.33, 131.76, 129.26, 128.98, 52.66, 52.19, 45.79, 36.56. MS (ESI+): 255.0 (M+H^+^).

**Dimethyl 2-(benzylthio)succinate**

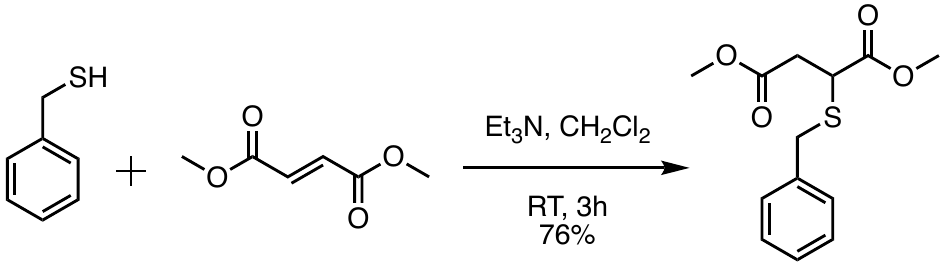

Dimethyl fumarate (0.173 g, 1.2 mmol) and triethylamine (0.167 mL, 1.2 mL) were added to a stirring solution of benzyl mercapatan (0.117 mL, 1.0 mmol) in dichloromethane (5 mL). The reaction mixture was stirred at room temperature and monitored by TLC. After 3 h, the reaction mixture was concentrated under reduced pressure and purified by flash chromatography (0🡪50% EtOAc:hexanes) to yield the product as white solid (76%). ^1^H NMR (400 MHz, CD_3_OD) δ 7.50-7.11 (m, 5H), 3.95-3.81 (m, 2H), 3.69 (s, 3H), 3.65-3.54 (m, 4H), 2.90 (dd, *J* = 17.0, 10.0 Hz, 1H), 2.60 (dd, *J* = 16.9, 5.6 Hz, 1H). ^13^C NMR (100 MHz, CD_3_OD) δ 173.75, 172.56, 138.74, 130.16, 129.54, 129.52, 128.31, 128.30, 52.86, 52.39, 42.22, 36.95, 36.80. MS (ESI+): 268.1 (M+H^+^).

**2-(Phenylthio)succinic acid**

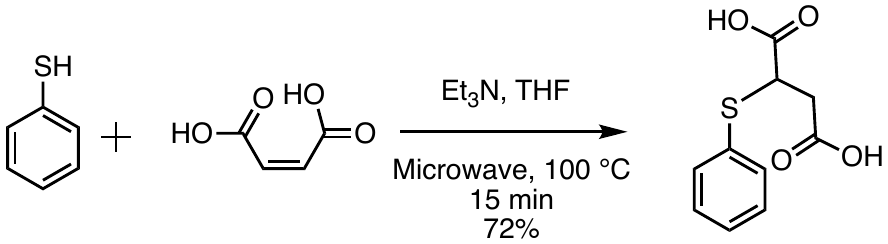

Maleic acid (0.139 g, 1.2 mmol), thiophenol (0.102 mL, 1.0 mmol) and triethylamine (0.167 mL, 1.2 mL) were dissolved in tetrahydrofuran (2 mL) and subjected to microwave irradiation in a 2-5 mL Biotage microwave vial at 100 °C for 15 minutes. The reaction mixture was concentrated under reduced pressure and was purified by flash chromatography (0🡪40% CH_3_OH:CH_2_Cl_2_) to yield the product as white solid (72%). ^1^H NMR (400 MHz, CD_3_OD) δ 7.58-7.47 (m, 2H), 7.41-7.28 (m, 3H), 3.95 (dd, *J* = 9.4, 5.7 Hz, 1H), 2.84 (dd, *J* = 17.0, 9.4 Hz, 1H), 2.69 (dd, *J* = 17.0, 5.7 Hz, 1H). ^13^C NMR (100 MHz, DMSO-*d*_6_) δ 172.00, 171.67, 132.51, 132.47, 129.14, 128.08, 45.02, 36.44. MS (ESI-): 225.0 (M-H^+^).

**2-(Benzylthio)succinic acid**

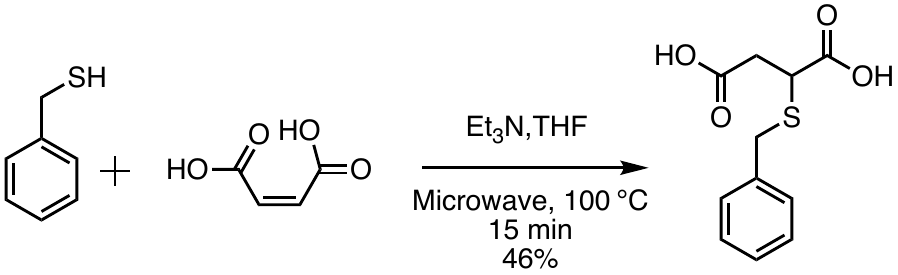

Maleic acid (0.139 g, 1.2 mmol), benzyl mercapatan (0.117 mL, 1.0 mmol) and triethylamine (0.167 mL, 1.2 mL) were dissolved in tetrahydrofuran (2 mL) and subjected to microwave irradiation in a 2-5 mL Biotage microwave vial at 100 °C for 15 minutes. The reaction mixture was concentrated under reduced pressure and was purified by flash chromatography (0🡪40% CH_3_OH:CH_2_Cl_2_) to yield the product as white solid (46%). ^1^H NMR (400 MHz, CD_3_OD) δ 7.57-7.01 (m, 5H), 4.00-3.82 (m, 2H), 3.53 (dd, *J* = 10.1, 5.1 Hz, 1H), 2.85 (dd, *J* = 17.0, 10.2 Hz, 1H), 2.54 (dd, *J* = 17.0, 5.1 Hz, 1H). ^13^C NMR (100 MHz, DMSO-*d*_6_) δ 172.61, 171.90, 137.61, 129.02, 128.53, 127.16, 48.66, 36.27, 34.97. MS (ESI-): 239.0 (M-H^+^).

**^1^H- and ^13^C-NMR spectra for FA-alkyne**

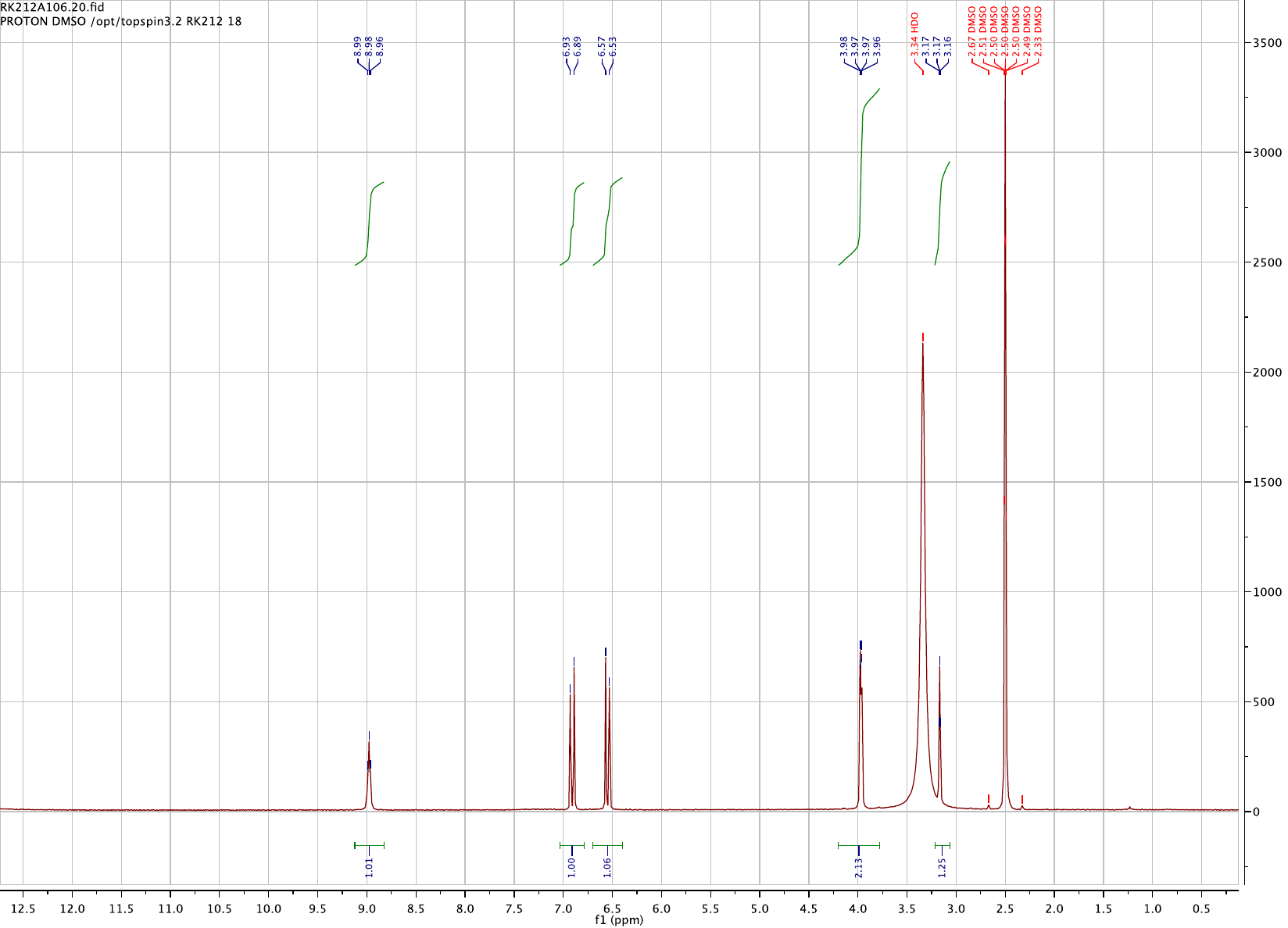

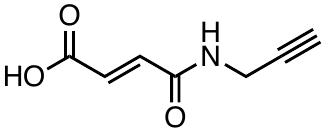

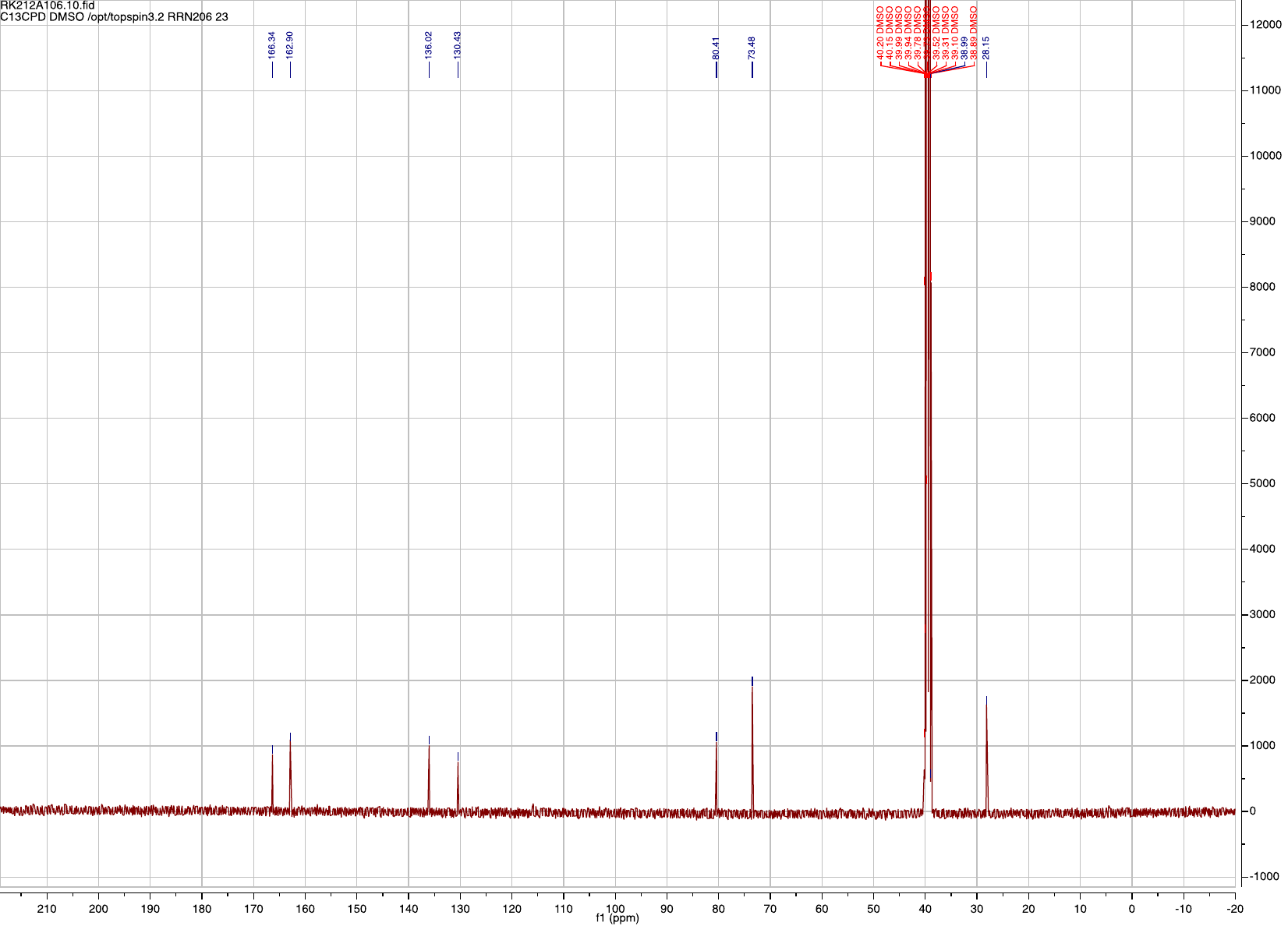

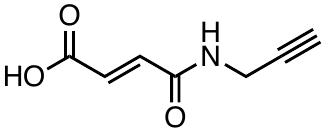

**^1^H- and ^13^C-NMR spectra for *S*-succinated thiols**

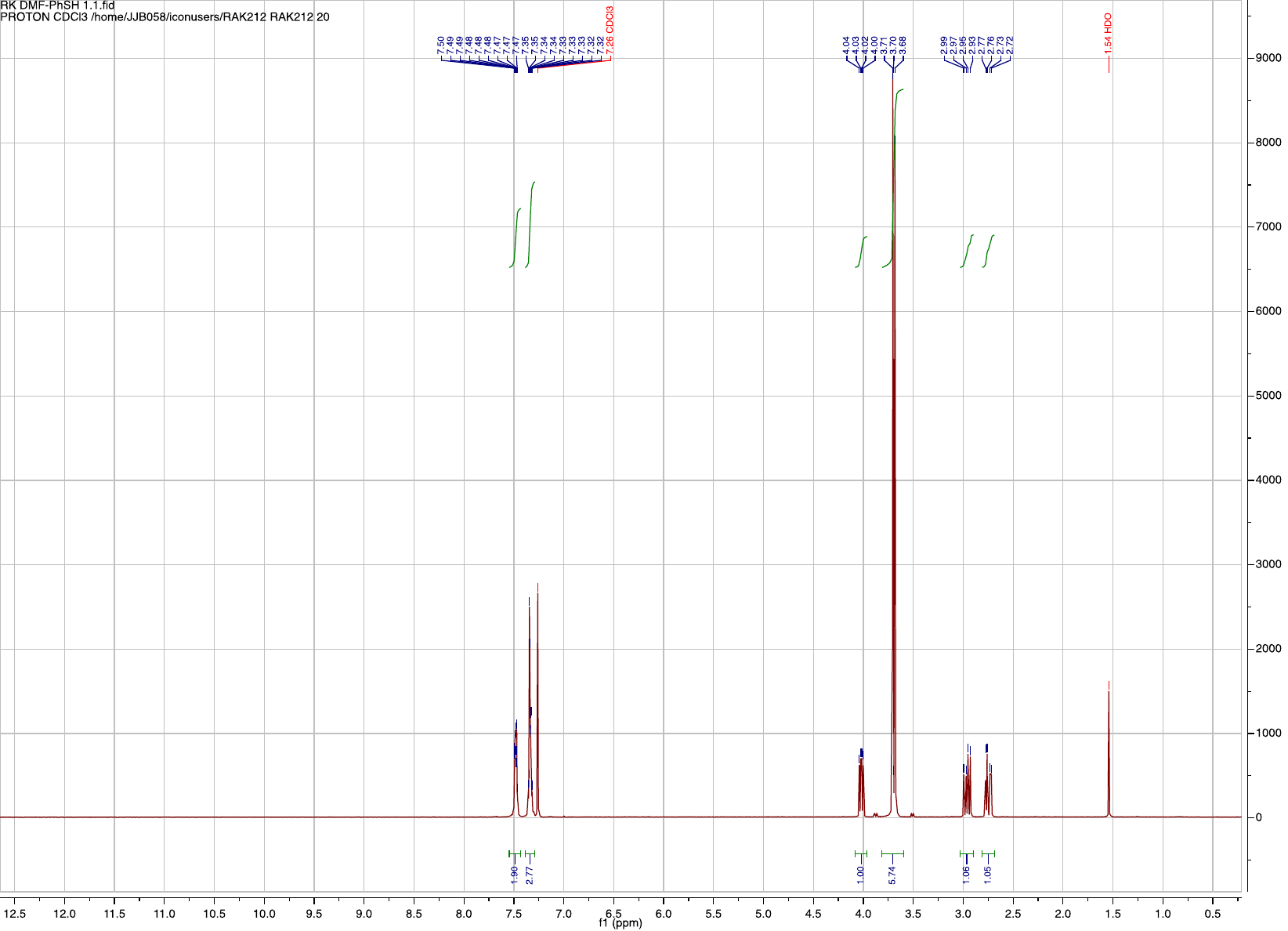

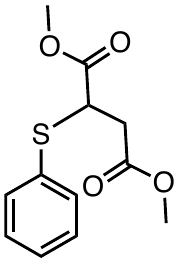

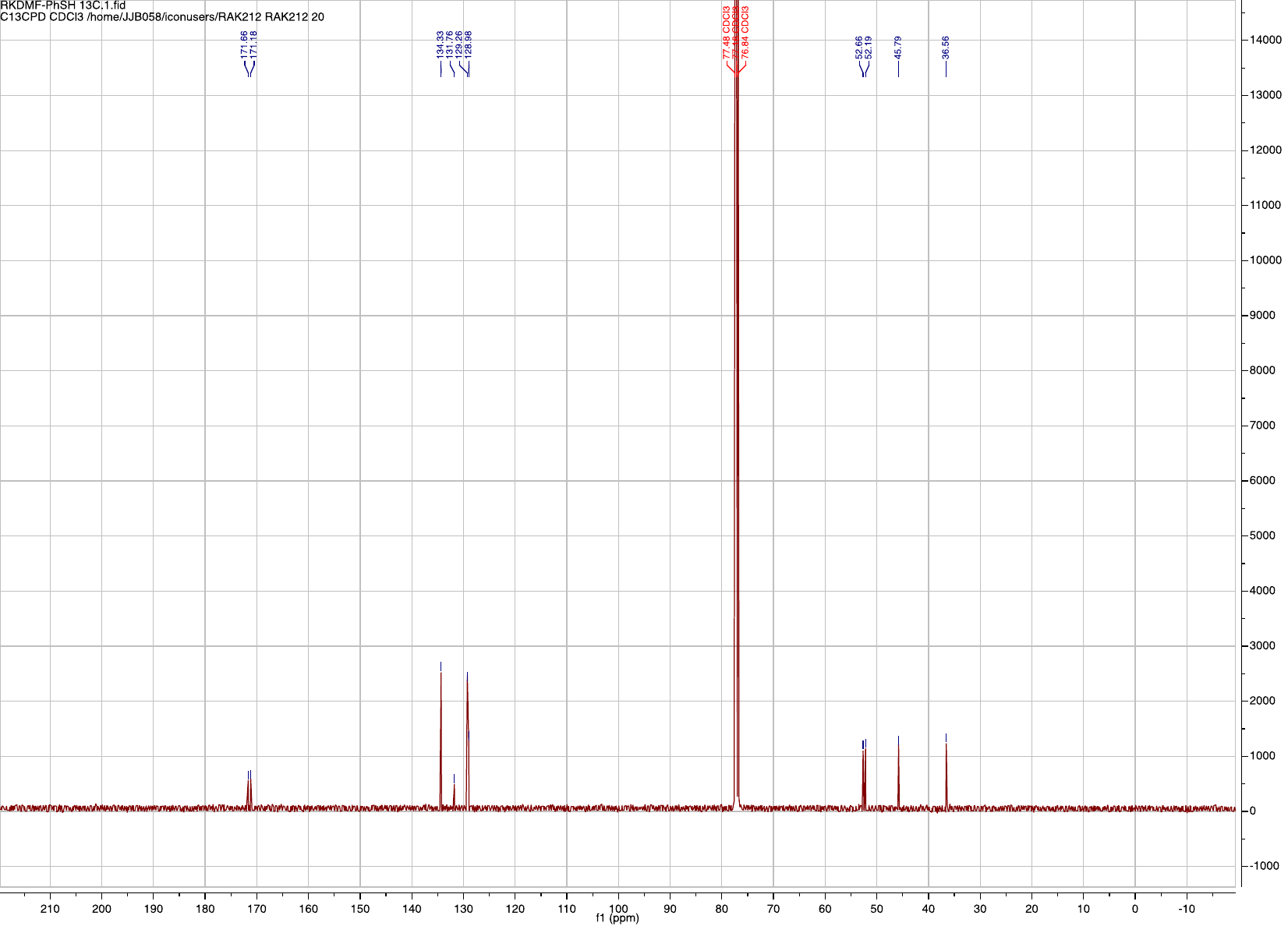

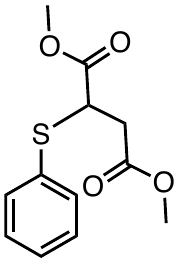

**^1^H- and ^13^C-NMR spectra for *S*-succinated thiols**

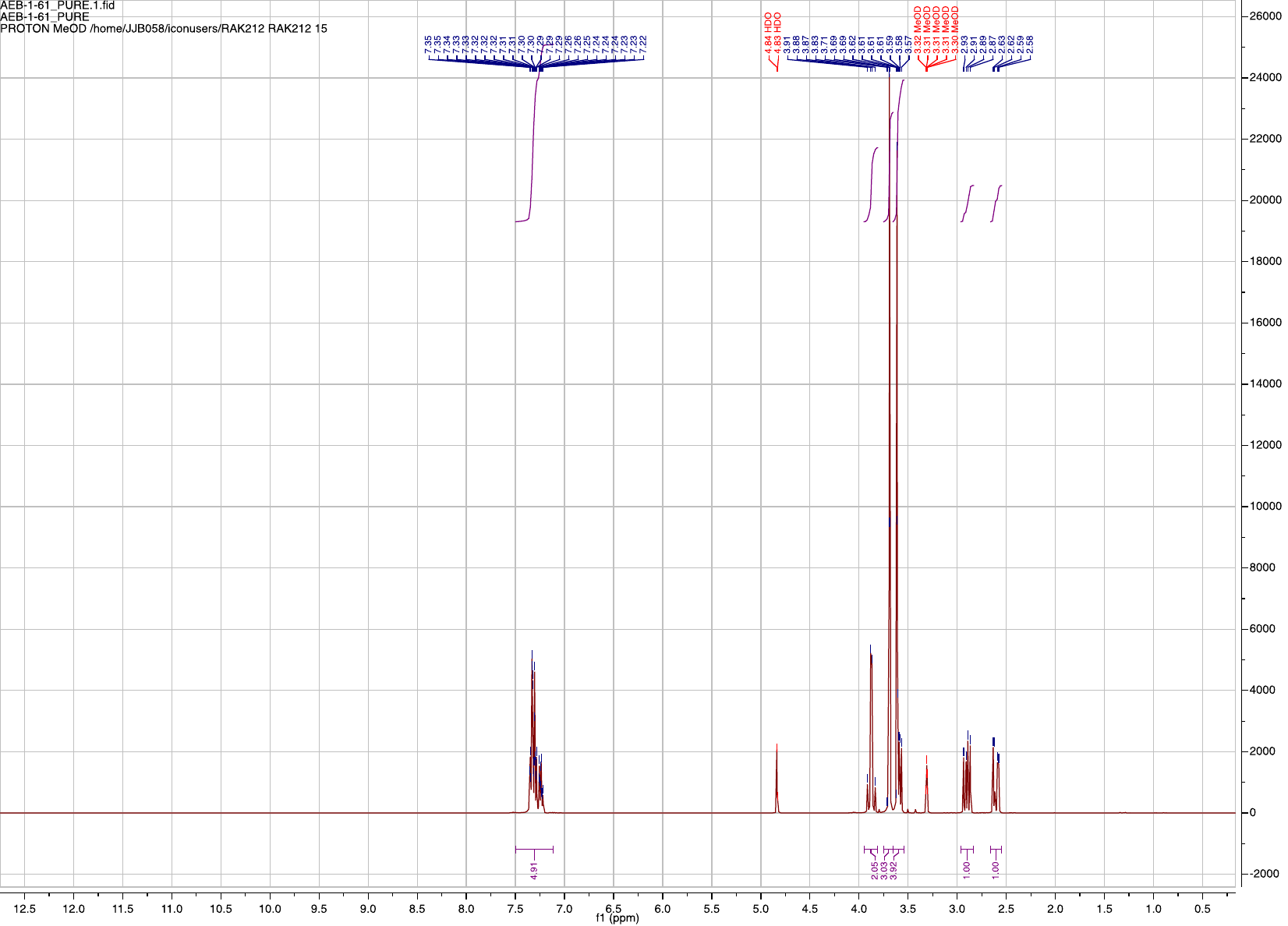

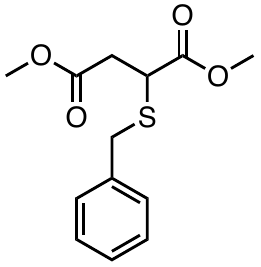

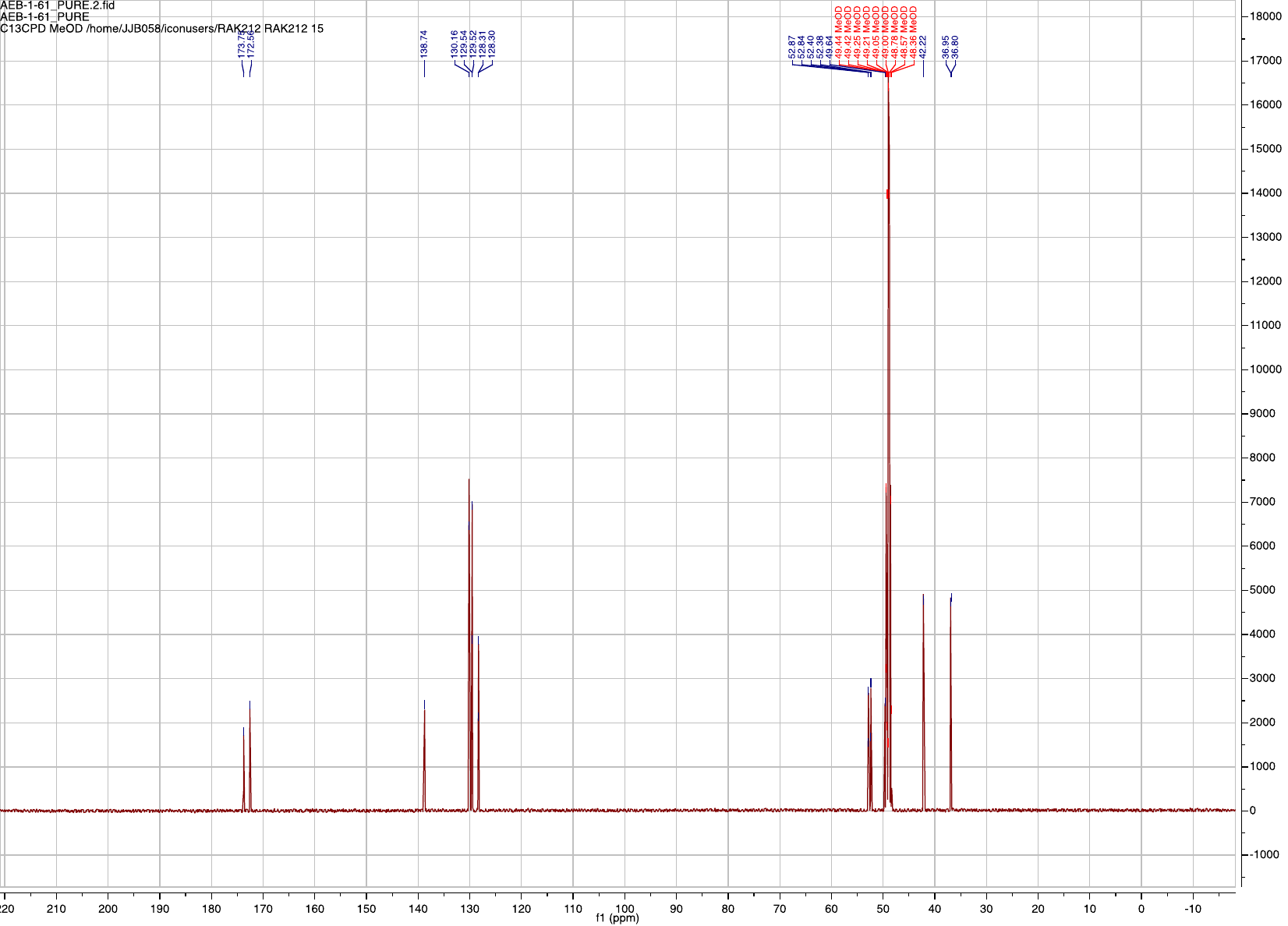

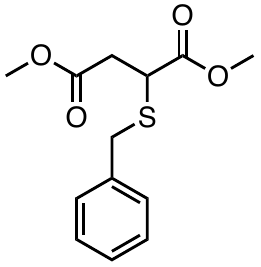
